# Computational Compensatory Mutation Discovery Approach: Predicting a PARP1 Variant Rescue Mutation

**DOI:** 10.1101/2021.11.21.469407

**Authors:** Krithika Ravishankar, Xianli Jiang, Emmett M. Leddin, Faruck Morcos, G. Andrés Cisneros

## Abstract

The prediction of protein mutations that affect function may be exploited for multiple uses. In the context of disease variants, the prediction of compensatory mutations that reestablish functional phenotypes could aid in the development of genetic therapies. In this work, we present an integrated approach that combines coevolutionary analysis and molecular dynamics (MD) simulations to discover functional compensatory mutations. This approach is employed to investigate possible rescue mutations of a poly(ADP-ribose) polymerase 1 (PARP1) variant, PARP1 V762A, associated with lung cancer and follicular lymphoma. MD simulations show PARP1 V762A exhibits noticeable changes in structural and dynamical behavior compared with wild type PARP1. Our integrated approach predicts A755E as a possible compensatory mutation based on coevolutionary information, and molecular simulations indicate that the PARP1 A755E/V762A double mutant exhibits similar structural and dynamical behavior to WT PARP1. Our methodology can be broadly applied to a large number of systems where single nucleotide polymorphisms (SNPs) have been identified as connected to disease and can shed light on the biophysical effects of such changes as well as provide a way to discover potential mutants that could restore wild type-like functionality. This can in turn be further utilized in the design of molecular therapeutics that aim to mimic such compensatory effect.

**Significance Statement:** Discovering protein mutations with desired phenotypes can be challenging due to its combinatorial nature. Herein we employ a methodology combining gene SNP association to disease, direct coupling analysis and molecular dynamics simulations to systematically predict rescue mutations. Our workflow identifies A755E as a potential rescue for the PARP1 V762A mutation, which has been associated with cancer. This methodology is general and can be applied broadly.

## Introduction

The identification of disease–associated mutations that result in missense protein variants can provide avenues for therapeutic development. For example, *trans*–splicing is a therapeutic approach that can be employed to repair mutations at the mRNA level (1). Therefore, understanding the impact of disease variants may be of value to determine if these mutations can or should be targeted for genetic therapies.

The identification and characterization of missense mutations has been a field of active research. Some of us have developed an approach termed Hypothesis Driven-SNP-Search (HyDn-SNP-S). This approach involves the search of single nucleotide polymorphisms (SNPs) resulting in missense mutations that are associated with a specific phenotype on a particular gene or genes, followed by atomistic simulations to characterize the impact of the mutation (2, 3). We have previously employed this approach to uncover and characterize various cancer– associated mutations (2, 4, 5), including the prediction and experimental confirmation of a rescue mutation for a lung–cancer associated mutation on APOBEC3H (6). Although there are successful examples for the prediction of rescue mutations, a systematic method to discover these variants would be beneficial.

Proteins evolve through a series of neutral or selectively-favored mutations (7, 8) that could coevolve with corresponding compensatory mutations to maintain constraints from folding structure or function (9–11). Such coevolutionary information between residue sites can be inferred by a statistical modeling of sequences in a protein family and has achieved significant performance in predicting physical contacts for protein folding and protein-protein interaction prediction (12–15). The coevolutionary model has also been used to estimate mutational effect in epistatis studies (16–19). The direct coupling analysis (DCA) method is a statistical model that estimates a global probability distribution of protein sequences by inferring parameters including covariation coupling between residues and site-wise conservation from multiple sequence alignments (MSAs) of homologous sequences (20). As a result, DCA is a useful tool that has been successfully applied in the prediction of protein structures (21, 22), conformational changes (23), protein interactions (24), function (25), Sequence Evolution with Epistatic Contributions (SEEC) (19), and recently in protein design (25, 26).

Given the features of these two methodologies, coevolutionary analysis and SNP search can be combined to further understand the relationship between cancer-related mutations and compensatory mutations which could rescue the SNP variant (Figure 2). Working from these two origins, molecular dynamics (MD) simulations of the identified mutations can be used to contextualize their impacts in reference to the wild type structure. In this contribution we present the development of a methodology that combines HyDn-SNP-S with coevolutionary analysis to uncover possible compensatory mutations for disease variants. We apply this approach to the regulatory domain of protein PARP1 and use MD simulations to understand the mutation’s effect on the overall PARP1 structure.

Poly(ADP-ribose) polymerase 1 (PARP1) performs base excision and repairs single-stranded breaks. It acts as an ADP-ribosylating enzyme, covalently attaching ADP-ribose to proteins. The successive transfer of ADP-ribose results a PAR chain, which acts as a signal for other DNA-repair enzymes (27). This process, known as PARsylation or PARylation, occurs on both single- and double-stranded DNA (28, 29). PARP1 is believed to perform over 90% of all cellular PARsylation activity (30). PARP1 is known to assist with the repair of single- and double-stranded breaks through several DNA repair pathways, including base excision repair (BER), nucleotide excision repair (NER), mismatch repair (MMR), homologous recombination repair (HRR), and non-homologous end joining (NHEJ) (28, 31). Despite its involvement in these various pathways, PARP1 is only essential for single-strand break repair, and it is considered non-essential for double-stranded repair (32). When it is in a position to repair strand breaks, PARP1 is believed to dimerize with the DNA-binding domain of another PARP1 (33). This dimerization is facilitated by the central automodification domain (34). PARP1 is of particular interest because of its inextricable link with the BRCA1 enzyme, known for breast cancer susceptibility (29, 35). Both PARP1 and BRCA1 are involved in homologous recombination repair (HRR), where a damaged area of DNA is resynthesized using a sister chromatid (36–38). BRCA1 is also known to perform PARsylation, which, together with RAP80, regulates HRR (39). This link has been utilized in the treatment of BRCA-mutated cancers, as evidenced by the FDA-approved use the PARP1 inhibitor Olaparib for advanced ovarian cancer (40). Inhibiting PARP1 leads to a stalling of the replication fork and the subsequent switch to repair via the NHEJ pathway in cancer cells, but a continuation via HRR in non-cancer cells (41).

PARP1’s N-terminal domain has three zinc fingers, one responsible for interactions between domains, and the other two involved in DNA binding (43). When DNA damage occurs, PARP1 localizes to the damaged area (44). The zinc fingers bind to the exposed nucleotides, instead of the 3’ and 5’ ends at the break sites, allowing for versatility in binding other secondary arrangements of DNA (45). The catalytic region then goes through three enzymatic reactions for PARsylation, composed of initiation, elongation, and branching. Central to this process is an “ADP-ribosyltransferase (ART) signature” (Figure 1) comprised of a conserved His-Tyr-Glu (H-Y-E) triad in its nicotinamide binding pocket (46). PARsylation requires the nicotinamide adenine dinucleotide (NAD^+^) as a coenzyme, because PARP1 polymerizes the ADP-ribose units (47). Unfolding of the helical subdomain (HD) (Figure 1) is crucial for the activation of PARP-1, and thus, changes in stability in this region can affect the enzyme’s catalytic output or how it binds NAD+ (48). This unfolding has been proposed to occur through a two-step mechanism, first through DNA binding and secondarily through substrate binding to destabilize the folded HD structure (49). Wild type PARP1 has been found to be upregulated in different cancers (50–56). In turn, overactivation of PARP1 can lead to mitochondrial distress and cell necrosis (57–59). One particular single nucleotide polymorphism (SNP), rs1136410, results in the V762A missense mutation in the HD region of PARP1. The resulting V762A mutation has been shown to reduce the enzymatic activity of PARP1 (60). This SNP has been linked to both lung cancer and follicular lymphoma through the HyDn-SNP-S method (2, 3). The rs1136410 SNP has also been shown to serve as a protective factor against breast cancer and coronary artery disease in the Han Chinese population, but it may lead to an increased overall risk of age-related cataracts and cancers (61–65).

**Figure 1:**
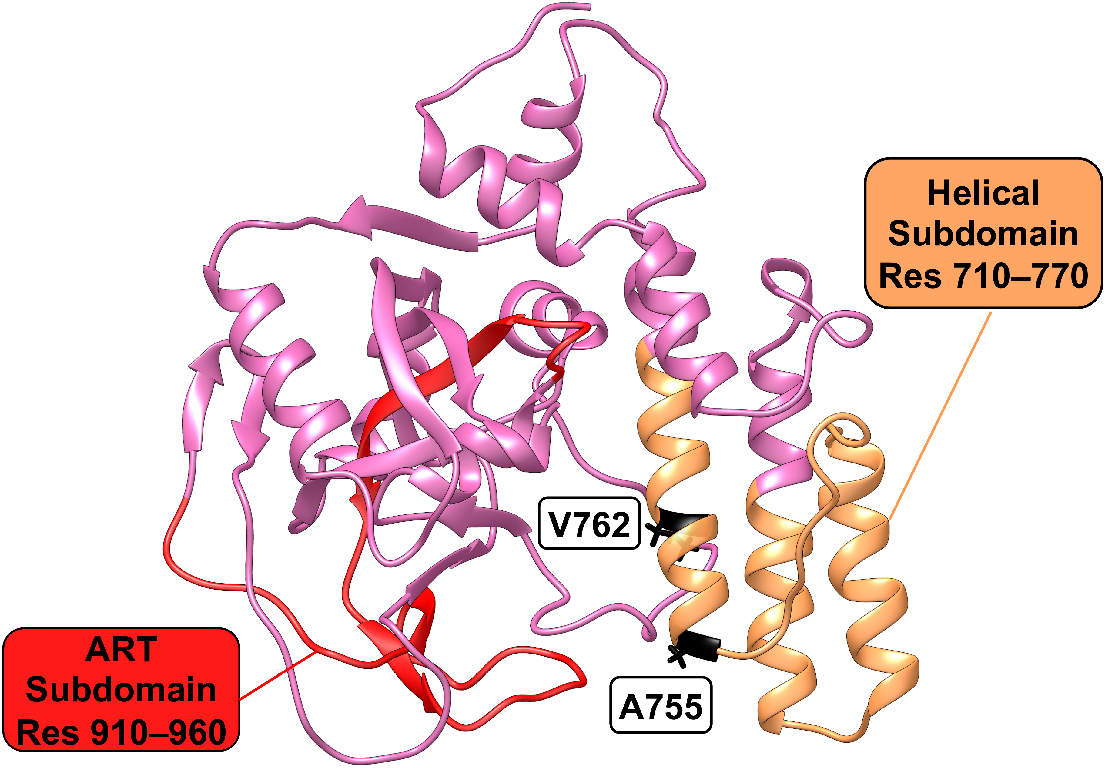
The ADP-ribosyltransferase (ART; red) and the helical (HD; pale orange) subdomains of the catalytic domain of PARP1 are highlighted on the 4ZZZ wild-type structure (42). Residues A755 and V762 are shown in black.

In the remainder of this paper we describe the application of the combined HyDn-SNP-S and coevolutionary analysis methods to determine whether there are possible compensatory mutations for the PARP1 V762A variant. In the next section we provide details of the coevolutionary analysis and molecular dynamics simulations methods, followed by the results of these approaches for wild type (WT) and various mutants of PARP1. Finally, concluding remarks are provided on the applicability of this combined approach.

## Materials and Methods

### Coevolutionary Analysis for PARP Regulatory Domain

The V762A mutation is found within the PARP regulatory domain of PARP1. To investigate evolutionary footprints for this functional domain, multiple sequence alignments (MSAs) of homologous sequences for this specific domain are obtained from the Pfam database with an entry ID of PF02877 (66). The direct coupling analysis (DCA) method (20) is then applied to the MSA dataset to extract information about coevolutionary coupling between any pairwise residues and the preference of amino acid occurrence at each residue position. As described in (20), DCA utilizes maximum entropy modeling to estimate the joint probability distribution of amino acid sequences of a protein or domain sequence:

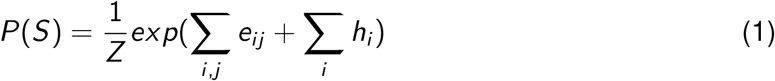

where *Z* is the partition function, the position of residues within the aligned domain or protein sequence are denoted as *i* and *j* and parameters *e_i,j_* and *h_i_* can be inferred by DCA. Parameters *e_i,j_* quantify coevolutionary coupling strength for residue *i* and *j* for all possible amino acid occurrence pairs. The amino acid biases for single residue positions is captured by the parameter *h_i_*. Being the exact inference of the parameters an intractable problem, there are multiple approximations to infer these parameters with different complexities and accuracies. In this work we use the mean field formulation (20), which optimizes the identification of highly coupled sites, however it is not generative as other approximations like bmDCA (67) or arDCA (68). Since the generative property is not useful in our context, mfDCA provides both accuracy and low computational complexity.

### Calculation of a Sequence-based Energy Function for PARP1 Mutants

Using the collection of *e_i,j_* and *h_i_* parameters estimated by DCA, a sequence-based energy function can be calculated from Equation 1 for any given aligned sequence. This collection of parameters or Hamiltonian (H) for a protein sequence S is expressed as:

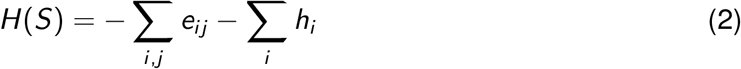

Calculating the energy function *H*(*S*_WT_) for the wild type sequence of PARP1’s regulatory domain provides a reference energy to compare against amino acid changes in the sequence. This sequence Hamiltoniann has been predictive of functional and non-functional effects in proteins and RNA (69–71). Any amino acid substitution in this domain would update the energy function to a mutant one *H*(*S*_Mut_). Then the effect of any mutant could be estimated in terms of the differential of this sequence-based energy function:

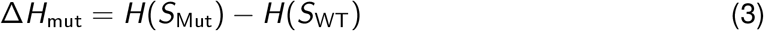

In this context, a more positive Δ*H*_mut_ score is predicted in general to have an unfavorable or neutral effect, while a more negative one represents a favorable change for fitness.

Original codes for coevolutionary parameter inference by DCA and H(S) score calculation were written in MATLAB and published before at https://github.com/morcoslab/coevolution-compatibility (72).

### Molecular Dynamics Analysis of PARP1

Seven systems of the catalytic domain in PARP1 were considered including: wild type from crystal 4ZZZ (WT) (42, 73), a cancer mutant containing the V762A mutation (rs1136410) from crystal 5WS1 (V762A) (74), a mutant containing the V762A mutation from the WT (V762A-from-WT), single mutants containing either A755E only (A755E) or A755L only (A755L), and double mutants containing the hypothesized rescue mutations and the V762A cancer-related mutation, A755E/V762A and A755L/V762A (Table 1). None of the systems studied contained DNA. The V762A-from-WT was created by using LEaP in AMBER Tools to edit structure 4ZZZ (75, 76). Modeller was used to incorporate the missing residues into both crystal structures (77, 78). The single mutants were created by editing structure 4ZZZ using UCSF Chimera and replacing the amino acid using the Dunbrack rotamer libraries (79, 80). The double mutants were similarly created by modifying structure 5WS1. VMD and UCSF Chimera were used for visualization (79, 81).

**Table 1:** Naming scheme for MD simulations of PARP1.

| Abbreviation | Mutations | Original PDB | Time Simulated (ns) |
| --- | --- | --- | --- |
| WT |  | 4ZZZ | 500 |
| V762A | V762A | 5WS1 | 500 |
| V762A-from-WT | V762A | 4ZZZ | 500 |
| A755E | A755E | 4ZZZ | 200 |
| A755L | A755L | 4ZZZ | 200 |
| A755E/V762A | A755E and V762A | 5WS1 | 200 |
| A755L/V762A | A755L and V762A | 5WS1 | 200 |

Using LEaP, chloride ions were added to neutralize the total charge of each system (75). WT, V762A, and V762A-from-WT were each solvated using TIP3P water extending at least 8 Å from the solute, and the A755E, A755L, A755E/V762A, and A755L/V762A systems were each solvated extending at least 12Å from the solute (75, 82). A simulation for a WT system with a 12 Å solvent buffer was also performed and no significant differences were observed compared with the smaller box results (Figures S4 and S7). Charged residues were assigned the default protonation state in LEaP, consistent with PROPKA (5 His had suggested protonation at N-delta by PROPKA, 3 of which were inconclusive by the electrostatic calculation and visual inspection) (83–85). The ff14SB force field was used for all protein residues (86).

AMBER molecular dynamics simulations were run using *pmemd.cuda* (76, 87, 88), with the NVT ensemble (number of atoms, volume, and temperature held constant) for the minimization and heating phases. The NPT ensemble (number of atoms, pressure, and temperature held constant) with the Langevin thermostat (temperature held at 300 K) was used for equilibration and production (89). The systems were run in triplicate with a 2 fs time step for the total simulation time shown in Table 1. Results for a representative trajectory of each system are shown below. All difference data between systems is presented as Variant - WT.

*Cpptraj* was used to analyze production dynamics (90). Normal modes were visualized using the Normal Mode Wizard in VMD (81, 91). Further data processing and graphing were performed with Gnuplot and the Matplotlib, NumPy, and statsmodels Python libraries (92–96). A FORTRAN90 program was used for the energy decomposition analysis (EDA) (97). EDA averaging was done using R (98), with the data.table, abind, and tidyverse libraries (99–101).

## Results and Discussion

We developed a compensatory mutation discovery workflow comprised of two computational approaches: 1) molecular dynamics simulations to investigate the effect of mutations on the protein’s structure and dynamics and 2) sequence-based coevolutionary analysis that provides a global single mutation landscape to screen out potential rescuing mutations for the SNP variant of interest. The compensatory mutation discovery workflow that has been developed herein is depicted schematically in Figure 2. Briefly, the two computational approaches are performed in tandem to investigate the structural and dynamic properties of the protein and disease variants under study via MD; coupled with the sequence-based coevolutionary analysis to obtain a single mutation landscape to screen out possible rescue mutations for the SNP variant of interest.

**Figure 2:**
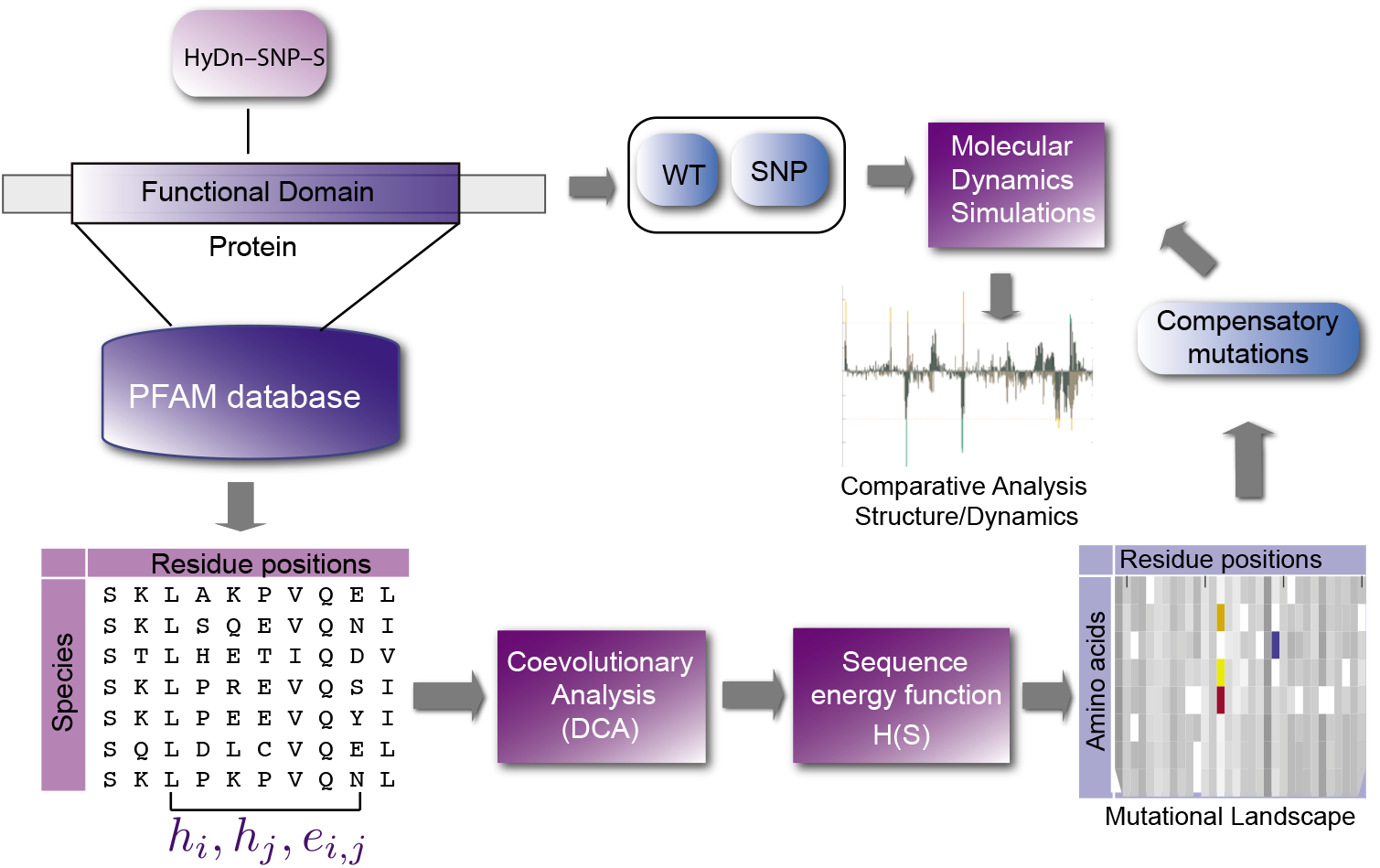
Workflow of DCA-MD method for identifying compensatory mutations for SNP(s). The DCA method infers coevolutionary parameter from MSA(s) containing the mutated residue (See Methods). Then a mutational landscape of protein energy function scores for all possible single mutations is generated to evaluate the SNP and initially screen possible compensatory mutations. The MD method simulates and validates the effect of SNP and compensatory mutation candidate.

Guided by the results from HyDn-SNP-S on PARP1, we sought to understand how the rs1136410 SNP affected the overall structure and dynamics of PARP1. Thus, we performed molecular dynamics (MD) simulations of both wild type PARP1 (WT) and the V762A PARP1 (V762A) variant structures.

Each system’s root mean square deviations (RMSDs) were stable across all simulations (see Figures S2–S3, and S8A). One way to assess the mutation’s effect on the dynamics of the system is through the use of a by-residue correlation matrix. This analysis can reveal regions of motion and dynamical correlation, anti-correlation, and no correlation within the protein (see Figures S9–S14). Based on the differences in correlated movements in Figure 3A, about half of the residues in the HD subdomain (710–770) and a fifth of the residues in the ART domain (910–960) show enhanced correlated movement in V762A than in WT.

**Figure 3:**
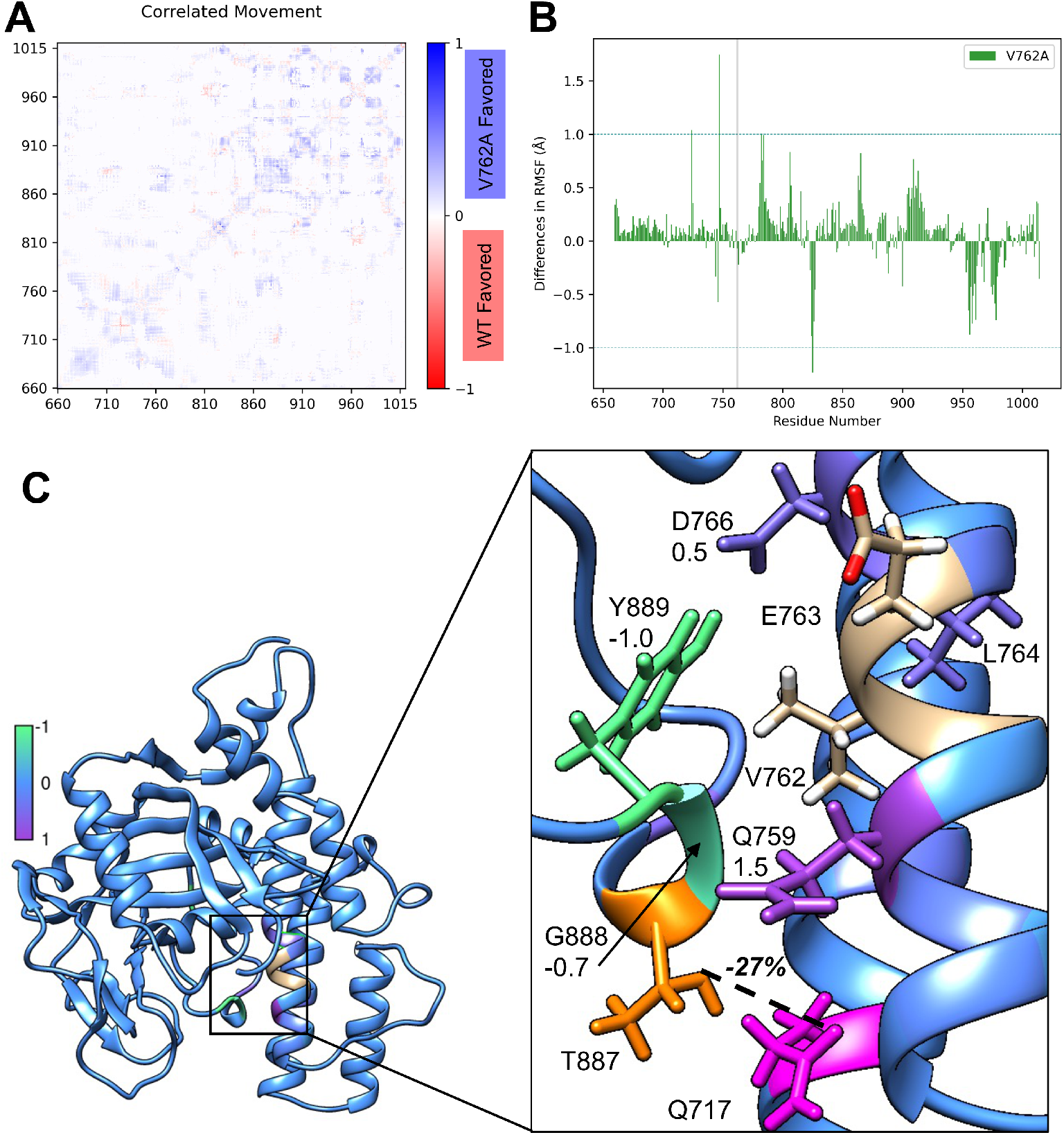
Comparison between V762A and WT. A. Differences in correlated movement between V762A (blue) and WT (red). B. Differences in RMSF between V762A and WT. A 1Å threshold is shown with a blue dashed line. C. Differences in EDA and hydrogen bonding between V762A and WT. Beige residues have undefined values. Differences in EDA above threshold of 0.5 Å are marked. Hydrogen bond donor in orange, acceptor in pink; bond indicated with bold dashed line.

An analysis of the root mean square fluctuation (RMSF) can be used to identify areas of higher or lower fluctuation between a system and its reference. Detailed RMSF data can be found in Table S1 and Figures S5–S7 and S8B,C. V762A and WT differ in RMSF by more than 1 Å at residues 724, 747, 782, and 825. Each of these residues is central to flexible loops throughout the subdomains, indicating a difference in dynamics between V762A and WT. V762A impacts the active site because of its proximity to the nicotinamide binding pocket. The active site residues (879 to 889; Figure 3B) show increased fluctuation in the V762A structure compared to WT, leading to decreased structural stability in the mutant. These residues are in a flexible loop opposite the NAD^+^-coordinating residues in the binding pocket, and several residues interact directly with V762.

An energy decomposition analysis (EDA), comprised of Coulomb and van der Waals (non-bonded) interactions, was used to study all of the intermolecular interactions between individual protein residues and residue 762 (see Figures S15–S25). Residues G888 and Y889, specifically, interact more favorably with residue 762 in the V762A mutant than in WT. The reverse behavior occurs with residue N759, which is located in the same helix as V762, but opposite the loop. N759 interacts more favorably with V762 in WT than in the V762A system (Figure 3C). Further, one of the hydrogen bonds between the HD and ART subdomains, GLN 717 – THR 887, is present for 27% less of production time in the V762A system than WT (Figure 3B). The V762A mutation appears to result in a reduction in stability in the active site because the loop and helix are not held as tightly together. This instability could mean that the NAD^+^ may bind with lower affinity in the PARP1 V762A holoenzyme.

We then utilized a DCA-based energy scoring function Δ*H*_Mut_ (see Materials and Methods) to explore the single mutation landscape for all residues in the regulatory domain of PARP1 (Figure 4A). The majority of single mutations had disruptive scores in for PAPR1 regulatory domain. V762A has a more positive score than WT, indicating its potential disruptive role in protein folding or stability from the perspective of coevolutionary analysis. Among all possible mutations, the top two most favorable mutations are observed in residue 755, specifically A755E and A755L. A double mutation profile generated with V762A also reports A755E and A755L as the best compensatory mutations occurring at positions other than 762 for the V762A mutant (Figure S1 and Figure 4B). This SNP-based profile directly estimates the effect of a second mutation on the original SNP variant, to uncover if there are second mutations that reverse the effect of SNP on the energy function score (Figure S1). Both A755E and A755L single mutations cause a comparable, but opposite, effect on protein coevolutionary score as V762A, while the double mutations, V762A/A755E and V762A/A755L, lead to scores near WT (Figure 4B,C). A755L generates a positive epistatic effect on V762A, suggesting that the double mutations has a better fitness energy score than the additive effect of two single mutations. A755E causes a negative epistatic effect on V762A. In summary, the coevolutionary analysis indicates that two mutations at residue 755 are promising for rescuing V762A.

**Figure 4:**
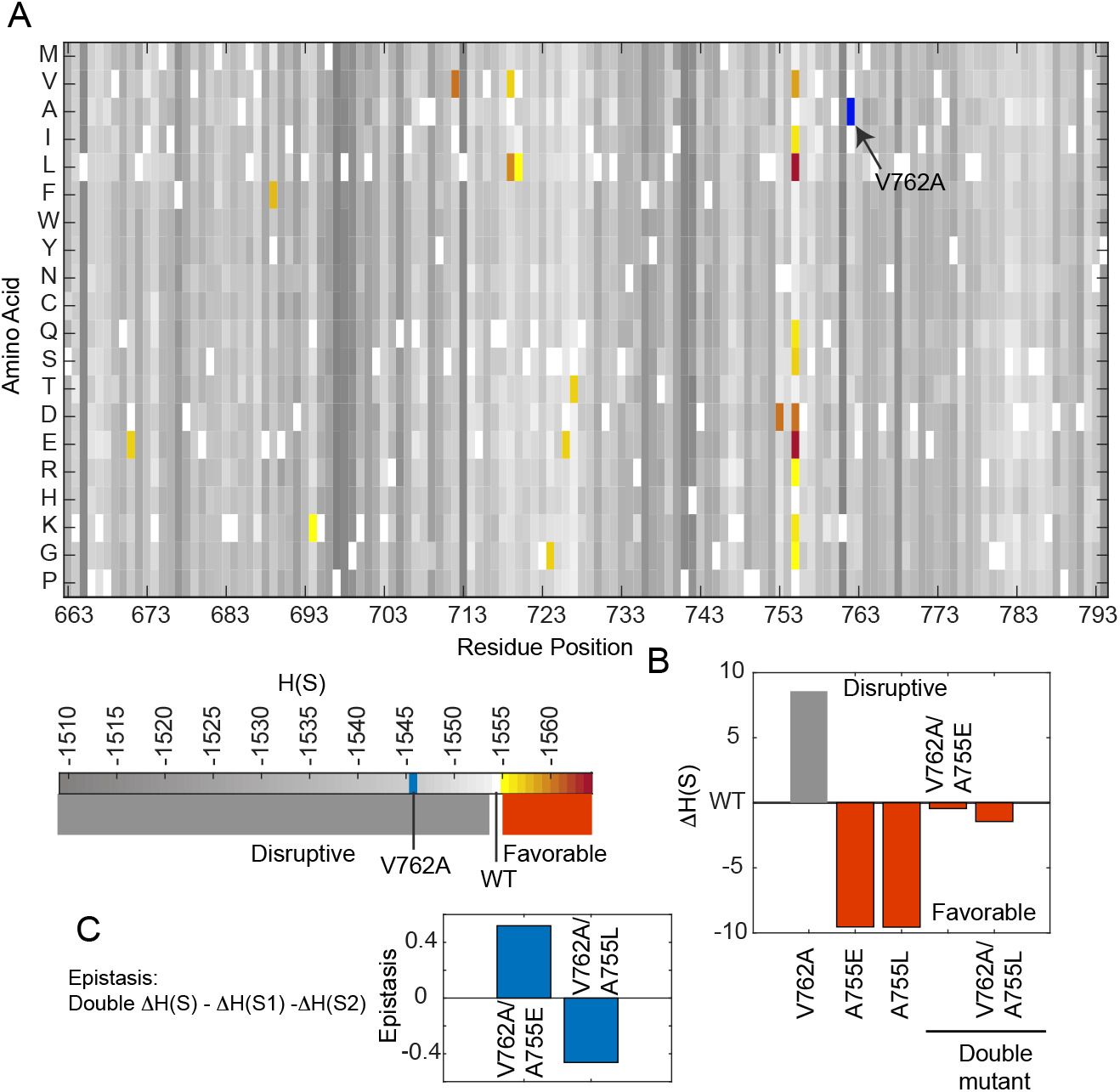
Coevolutionary information based energy for PARP1 and mutant. A. Energy landscape of mutations on PARP regulatory domain for PARP1. B. Mutational effect of PARP1 SNP V762A and potential complementary mutants, A755E/L. C. Epistatic effect for V755E and V755L for PARP1 SNP V762A.

Working from the results of the coevolutionary analysis, we simulated the A775E and A755L single mutants to establish a baseline for those mutations. The hydrogen bond between GLN 717 and THR 887 is present 28% (OE1–OG1) and 23% (OE1–N) less of production time in the A755L system than in the WT, indicating that A755L leads to less stability in the active site (lower box of Figure 5A). The EDA revealed significant differences in the non-bonded interactions between A755E and WT; with a large number of residues in the catalytic domain showing changes larger than 1 kcal/mol (Figure 5B). This may be due to the change from a residue with no charge to one that is negatively charged. Additionally, several HD subdomain residues (717, 720, 758–759) and residue 887, which is located in the active site, all interact more favorably with residue 755 in A755L than in WT (Figure 5B). Four residues in flexible loops, two in the HD subdomain (744 and 746) and two in the ART subdomain (824 and 825), have a significantly lower RMSF in A755E and A755L than WT (Figure 5C and S8C), suggesting that both A755E and A755L stabilize the overall structure as a result of this decreased flexibility. Based on the differences in correlated movements, the portion of the helices of the HD subdomain near the variant (residues 710–770) show more correlated movement in both A755E and A755L than in WT with themselves (Figure 5D-E).

**Figure 5:**
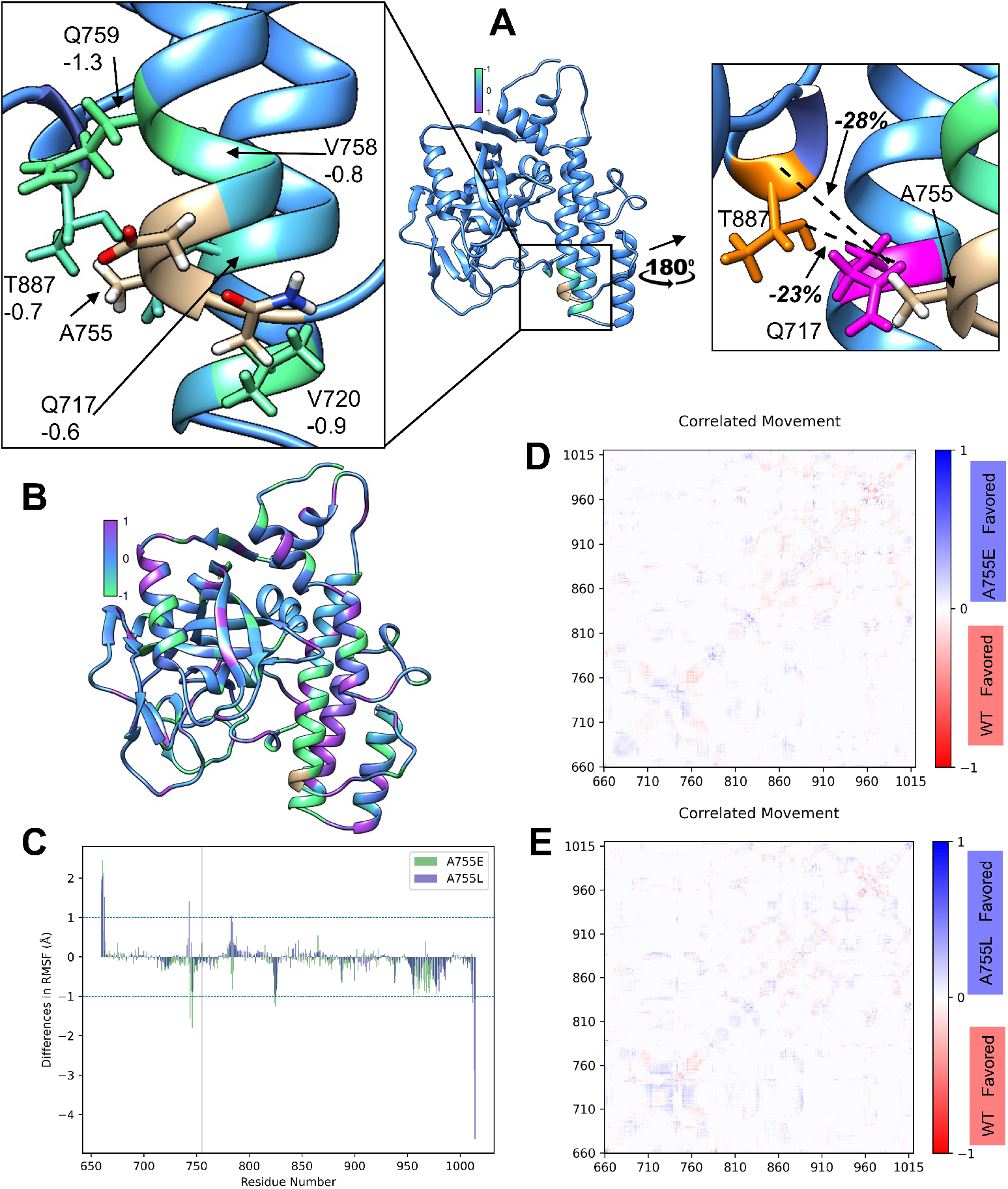
Comparison between A755E and WT and A755L and WT. A. Differences in EDA and hydrogen bonding between A755L and WT. Beige residues have undefined values. Differences in EDA above threshold of 0.5 kcal/mol are marked. Hydrogen bond donor in orange, acceptor in pink; bonds indicated with bold dashed line. B. Differences in EDA between A755E and WT. Beige residues have undefined values. Differences in EDA above threshold of 0.5 kcal/mol are marked. C. Differences in RMSF between A755E and A755L and WT. A 1 Å threshold is shown with a blue dashed line. D. Differences in correlated movement between A755E (blue) and WT (red). E. Differences in correlated movement between A755L (blue) and WT (red).

We then simulated the A755E/V762A and A755L/V762A double mutant systems to evaluate the role of the predicted residues as compensatory mutations on the dynamics of PARP1. The hydrogen bond between GLN 717 and THR 887 is present 28% (OE1–OG1) less of production time in A755L/V762A than in the WT, indicating that A755L/V762A leads to less stability in the active site (Figure 6A). Residues 717, 720, 758–759, which are near the site of mutation, and 887, which is located in the active site, all interact more favorably with residue 762 in A755L/V762A than in WT (Figure 6A).

**Figure 6:**
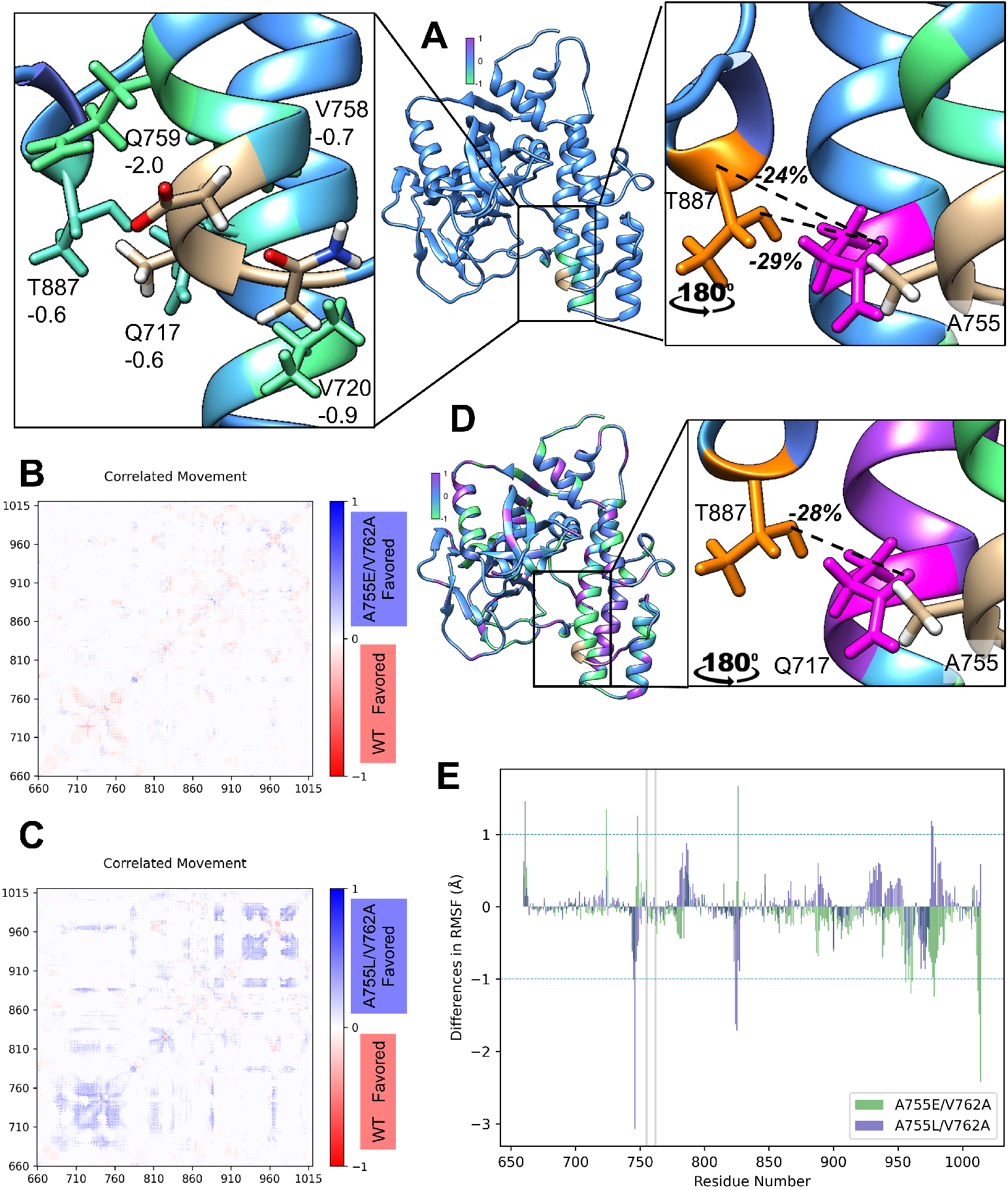
Comparison between A755E/V762A and WT and A755L/V762A and WT. A. Differences in EDA and hydrogen bonding between A755L/V762A and WT. Beige residues have undefined values. Differences in EDA above threshold of 0.5kcal/mol are marked. Hydrogen bond donor in orange, acceptor in pink; bonds indicated with bold dashed line. B. Differences in correlated movement between A755E/V762A (blue) and WT (red). C. Differences in correlated movement between A755L/V762A (blue) and WT (red). D. Differences in EDA and hydrogen bonding between A755E/V762A and WT. Beige residues have undefined values. Differences in EDA above threshold of 0.5kcal/mol are marked. Hydrogen bond donor in orange, acceptor in pink; bonds indicated with bold dashed line. E. Differences in RMSF between A755E/V762A and A755L/V762A and WT. A 1Å threshold is shown with a blue dashed line.

There is minimal difference in correlated movements between A755E/V762A and WT, potentially indicating that A755E is a rescue mutation (Figure 6B). Based on the differences in correlated movements, residues 710-770 show more correlated movement in A755L/V762A than in WT with themselves and 885-985 with themselves (Figure 6C). This impact on the HD subdomain may point to a similar or increased catalytic output for structures with A755E/L rescue mutations.

In A755E/V762A, the hydrogen bond between GLN 717 and THR 887 is present 29% (OE1–OG1) and 24% (OE1–N) less of production time than in the WT, indicating that A755E/V762A also leads to less stability in the active site (lower box of Figure 6D). There are significant differences in the non-bonded interactions between A755E/V762A and WT; residues all over the catalytic domain are impacted (Figure 6D). Similar to A755E, the changes may be due to the additional charge at position 755.

At residues 724, 748, and 826, the RMSF was significantly lower in A755E/V762A than WT (Figure 6E). At residues 960, 961, and 968, the RMSF was significantly higher in A755E/V762A than WT (Figure 6E). These correspond to differences seen for A755E/V762A in the normal modes analysis (see Figures S26–S27), where the HD subdomain shows less motion than the ART subdomain. The A755L/V762A system, however, closely resembles the WT in its first normal mode. As these residues are indicated by RMSF in each mutant studied, their fluctuation may be important for the recognition of the cofactor, which is absent from these simulations. The added stability provided by A755L in the A755L/V762A gives strong support for its evolution as a compensatory mutation. Because HD region destabilization is necessary for PARP1 activation, this particular double-mutant may more tightly control activation.

## Conclusion

In this study, we have demonstrated that integrating coevolutionary analysis and MD simulations can be useful to discover and validate compensatory mutations for SNPs using PARP1 rs1136410 as an example. A755E/L is first recognized by the DCA coevolutionary method as variants that are most favorable for PARP1 structures and the subsequent MD simulations validated that both variants stabilize the overall structure. Coevolutionary information can also be used to estimate double mutations that contain SNP to uncover rescue mutations. Both A755E/L lead to favorable “fitness” conditions in the context of the V762A variant from an evolutionary perspective. Additionally, the effects of A755E/L and V762A on PARP1 protein are not purely additive, with A755E being negatively synergetic and A755L being positively synergetic (Figure 4B). MD simulations show that the cancer mutation affects the structure and dynamics of V762A PARP1 compared with WT. These results indicate that the A775E mutation, in conjunction with V762A, can resolve some of the structural and dynamical impacts, mimicking wild type. Our work can help understand the effects of SNPs in their association with disease, like cancer in this case, as well as identifying changes that could ameliorate those changes. The discovery of important compensatory mutations can be used to study how particular SNPs are not always associated with disease and provide a roadmap for molecular therapeutic approaches aiming at reducing the negative effects of mutations. This methodology is generic in the sense that can be applied to a large number of systems where structural and sequence data is available. The case of PARP1, presented here, is only one of many that could be studied with our integrated approach. Subsequent computational and new experimental investigation of the potential of the two proposed rescue mutations would provide further insights. We expect future work could uncover important insights on the effect of mutations for many more genes and their associated diseases.

## Author Contributions

KR, XJ, FM, and GAC designed the project. KR and EML carried out MD simulations, analyzed and interpreted data, and co-wrote the manuscript. XJ performed coevolutionary analysis, analyzed data and co-wrote the manuscript. All authors contributed to discussion and manuscript editing.

## Acknowledgments

This work was supported by NIH R01GM108583 (GAC), R35GM133631 (XJ and FM), and NSF CAREER grant MCB-1943442 (FM). Computational time from CASCaM’s CRUNTCH3 cluster, partially supported by NSF CHE-1531468 (GAC), and from XSEDE Project No. TG-CHE160044 is thankfully acknowledged.

## References

1. Hong, E. M., C. K. Ingemarsdotter, and A. M. L. Lever, 2020. Therapeutic applications of trans-splicing. Br. Med. Bull. 136:4–20, DOI: 10.1093/bmb/ldaa028.

2. Swett, R. J., A. Elias, J. A. Miller, G. E. Dyson, and G. Andrés Cisneros, 2013. Hypothesis driven single nucleotide polymorphism search (HyDn-SNP-S). DNA Repair 12:733–740, DOI: 10.1016/j.dnarep.2013.06.001.

3. Silvestrov, P., S. J. Maier, M. Fang, and G. A. Cisneros, 2018. DNArCdb: A database of cancer biomarkers in DNA repair genes that includes variants related to multiple cancer phenotypes. DNA Repair 70:10–17, DOI: 10.1016/j.dnarep.2018.07.010.

4. Walker, A. R., P. Silvestrov, T. A. Müller, R. H. Podolsky, G. Dyson, R. P. Hausinger, and G. A. Cisneros, 2017. ALKBH7 variant related to prostate cancer exhibits altered substrate binding. PLOS Computational Biology 13:1–13, DOI: 10.1371/journal.pcbi.1005345.

5. Antczak, N. M., A. R. Walker, H. R. Stern, E. M. Leddin, C. Palad, T. A. Coulther, R. J. Swett, G. A. Cisneros, and P. J. Beuning, 2018. Characterization of nine cancer-associated variants in human DNA polymerase *»*. Chem. Res. Toxicol. 31:697–711, DOI: 10.1021/acs.chemrestox.8b00055.

6. Hix, M. A., L. Wong, B. Flath, L. Chelico, and G. A. Cisneros, 2020. Singlenucleotide polymorphism of the DNA cytosine deaminase APOBEC3H haplotype I leads to enzyme destabilization and correlates with lung cancer. NAR Cancer 2:zcaa023, DOI: 10.1093/narcan/zcaa023.

7. Camps, M., A. Herman, E. Loh, and L. A. Loeb, 2007. Genetic constraints on protein evolution. Crit. Rev. Biochem. Mol. Biol. 42:313–326, DOI: 10.1080/10409230701597642.

8. Storz, J. F., 2016. Causes of molecular convergence and parallelism in protein evolution. Nat. Rev. Genet. 17:239–250, DOI: 10.1038/nrg.2016.11.

9. Davis, B. H., A. F. Poon, and M. C. Whitlock, 2009. Compensatory mutations are repeatable and clustered within proteins. Proc. R. Soc. B 276:1823–1827, DOI: 10.1098/rspb.2008.1846.

10. Saikat Chakrabarti, A. R. P., 2010. Structural and functional roles of coevolved sites in proteins. PLoS ONE 5:e8591, DOI: 10.1371/journal.pone.0008591.

11. de Juan, D., F. Pazos, and A. Valencia, 2013. Emerging methods in protein co-evolution. Nat. Rev. Genet. 14:249–261, DOI: 10.1038/nrg3414.

12. Morcos, F., and J. N. Onuchic, 2019. The role of coevolutionary signatures in protein interaction dynamics, complex inference, molecular recognition, and mutational landscapes. Curr. Opin. Struct. Biol. 56:179–186, DOI: 10.1016/j.sbi.2019.03.024.

13. Jones, D. T., D. W. A. Buchan, D. Cozzetto, and M. Pontil, 2011. PSICOV: Precise structural contact prediction using sparse inverse covariance estimation on large multiple sequence alignments. Bioinformatics 28:184–190, DOI: 10.1093/bioinformatics/btr638.

14. Kamisetty, H., S. Ovchinnikov, and D. Baker, 2013. Assessing the utility of coevolution-based residue–residue contact predictions in a sequence-and structure-rich era. Proc. Natl. Acad. Sci. U. S. A. 110:15674–15679, DOI: 10.1073/pnas.1314045110.

15. Marks, D. S., L. J. Colwell, R. Sheridan, T. A. Hopf, A. Pagnani, R. Zecchina, and C. Sander, 2011. Protein 3D structure computed from evolutionary sequence variation. PLOS ONE 6:1–20, DOI: 10.1371/journal.pone.0028766.

16. Flynn, W. F., A. Haldane, B. E. Torbett, and R. M. Levy, 2017. Inference of Epistatic Effects Leading to Entrenchment and Drug Resistance in HIV-1 Protease. Molecular Biology and Evolution 34:1291–1306, DOI: 10.1093/molbev/msx095.

17. Biswas, A., A. Haldane, E. Arnold, and R. M. Levy, 2019. Epistasis and entrenchment of drug resistance in HIV-1 subtype B. eLife 8, DOI: 10.7554/eLife.50524.

18. Bisardi, M., J. Rodriguez-Rivas, F. Zamponi, and M. Weigt, 2021. Modeling sequence-space exploration and emergence of epistatic signals in protein evolution. Mol. Biol. Evol. 39, DOI: 10.1093/molbev/msab321.

19. de la Paz, J. A., C. M. Nartey, M. Yuvaraj, and M. Faruck, 2020. Epistatic contributions promote the unification of incompatible models of neutral molecular evolution. Proc. Natl. Acad. Sci. U. S. A. 117, DOI: 10.1073/pnas.1913071117.

20. Morcos, F., A. Pagnani, B. Lunt, A. Bertolino, D. S. Marks, C. Sander, R. Zecchina, J. N. Onuchic, T. Hwa, and M. Weigt, 2011. Direct-coupling analysis of residue coevolution captures native contacts across many protein families. Proc. Natl. Acad. Sci. U. S. A. 108, DOI: 10.1073/pnas.1111471108.

21. dos Santos, R., X. Jiang, L. Martínez, and M. F, 2019. Coevolutionary signals and structure-based models for the prediction of protein native conformations. Computational Methods in Protein Evolution 1851, DOI: 10.1007/978-1-4939-8736-8_5.

22. Dos Santos, R., A. Ferrari, H. de Jesus, F. Gozzo, F. Morcos, and L. Martinez, 2018. Enhancing protein fold determination by exploring the complementary information of chemical cross-linking and coevolutionary signals. Bioinformatics 34, DOI: 10.1093/bioinformatics/bty074.

23. Morcos, F., B. Jana, T. Hwa, and J. Onuchic, 2013. Coevolutionary signals across protein lineages help capture multiple protein conformations. Proc. Natl. Acad. Sci. U. S. A. 110, DOI: 10.1073/pnas.1315625110.

24. dos Santos, R., F. Morcos, B. Jana, A. Andricopulo, and J. Onuchic, 2015. Dimeric interactions and complex formation using direct coevolutionary couplings. Sci. Rep. 5, DOI: 10.1038/srep13652.

25. Dimas, R. P., X.-L. Jiang, J. Alberto de la Paz, F. Morcos, and C. T. Chan, 2019. Engineering repressors with coevolutionary cues facilitates toggle switches with a master reset. Nucleic Acids Res. 47:5449–5463, DOI: 10.1093/nar/gkz280.

26. Jiang, X. L., R. P. Dimas, C. T. Y. Chan, and F. Morcos, 2021. Coevolutionary methods enable robust design of modular repressors by reestablishing intra-protein interactions. Nat Commun. 12:5592, DOI: 10.1038/s41467-021-25851-6.

27. Wei, H., and X. Yu, 2016. Functions of PARylation in DNA damage repair pathways. Genomics, Proteomics Bioinf. 14:131–139, DOI: 10.1016/j.gpb.2016.05.001.

28. Ko, H. L., and E. C. Ren, 2012. Functional aspects of PARP1 in DNA repair and transcription. Biomolecules 2:524–548, DOI: 10.3390/biom2040524.

29. Hu, Y., S. A. Petit, S. B. Ficarro, K. J. Toomire, A. Xie, E. Lim, S. A. Cao, E. Park, M. J. Eck, R. Scully, M. Brown, J. A. Marto, and D. M. Livingston, 2014. PARP1-driven poly-ADP-ribosylation regulates BRCA1 function in homologous recombination–mediated DNA repair. Cancer Discovery 4:1430–1447, DOI: 10.1158/2159-8290.CD-13-0891.

30. Kamaletdinova, T., Z. Fanaei-Kahrani, and Z.-Q. Wang, 2019. The enigmatic function of PARP1: From PARylation activity to PAR readers. Cells 8, DOI: 10.3390/cells8121625.

31. Wang, M., W. Wu, W. Wu, B. Rosidi, L. Zhang, H. Wang, and G. Iliakis, 2006. PARP-1 and Ku compete for repair of DNA double strand breaks by distinct NHEJ pathways. Nucleic Acids Res. 34:6170–6182, DOI: 10.1093/nar/gkl840.

32. Ray Chaudhuri, A., and A. Nussenzweig, 2017. The multifaceted roles of PARP1 in DNA repair and chromatin remodelling. Nat. Rev. Mol. Cell Biol. 18:610–621, DOI: 10.1038/nrm.2017.53.

33. Ali, A. A. E., G. Timinszky, R. Arribas-Bosacoma, M. Kozlowski, P. O. Hassa, M. Hassler, A. G. Ladurner, L. H. Pearl, and A. W. Oliver, 2012. The zinc-finger domains of PARP1 cooperate to recognize DNA strand breaks. Nat. Struct. Mol. Biol. 19:685–692, DOI: 10.1038/nsmb.2335.

34. Mendoza-Alvarez, H., and R. Alvarez-Gonzalez, 1993. Poly(ADP-ribose) polymerase is a catalytic dimer and the automodification reaction is intermolecular. J. Biol. Chem. 268:22575–22580, DOI: 10.1016/S0021-9258(18)41568-2.

35. Li, D., F.-F. Bi, N.-N. Chen, J.-M. Cao, W.-P. Sun, Y.-M. Zhou, C.-Y. Li, and Q. Yang, 2014. A novel crosstalk between BRCA1 and poly (ADP-ribose) polymerase 1 in breast cancer. Cell Cycle 13:3442–3449, DOI: 10.4161/15384101.2014.956507.

36. Li, X., and W.-D. Heyer, 2008. Homologous recombination in DNA repair and DNA damage tolerance. Cell Res. 18:99–113, DOI: 10.1038/cr.2008.1.

37. Prakash, R., Y. Zhang, W. Feng, and M. Jasin, 2015. Homologous recombination and human health: The roles of BRCA1, BRCA2, and associated proteins. Cold Spring Harbor Perspect. Biol. 7, DOI: 10.1101/cshperspect.a016600.

38. Li, H., Z.-Y. Liu, N. Wu, Y.-C. Chen, Q. Cheng, and J. Wang, 2020. PARP inhibitor resistance: the underlying mechanisms and clinical implications. Mol. Cancer 19:107, DOI: 10.1186/s12943-020-01227-0.

39. Vohhodina, J., K. J. Toomire, S. A. Petit, G. Micevic, G. Kumari, V. V. Botchkarev, Z. Li, D. M. Livingston, and Y. Hu, 2020. RAP80 and BRCA1 PARsylation protect chromosome integrity by preventing retention of BRCA1-B/C complexes in DNA repair foci. Proc. Natl. Acad. Sci. U. S. A. 117:2084–2091, DOI: 10.1073/pnas.1908003117.

40. Kim, G., G. Ison, A. E. McKee, H. Zhang, S. Tang, T. Gwise, R. Sridhara, E. Lee, A. Tzou, R. Philip, H.-J. Chiu, T. K. Ricks, T. Palmby, A. M. Russell, G. Ladouceur, E. Pfuma, H. Li, L. Zhao, Q. Liu, R. Venugopal, A. Ibrahim, and R. Pazdur, 2015. FDA approval summary: Olaparib monotherapy in patients with deleterious germline BRCA-mutated advanced ovarian cancer treated with three or more Lines of chemotherapy. Clin. Cancer Res. 21:4257–4261, DOI: 10.1158/1078-0432.CCR-15-0887.

41. Pilie, P., C. M. Gay, L. A. Byers, M. J. O’Connor, and T. A. Yap, 2019. PARP inhibitors: extending benefit beyond BRCA mutant cancers. Clin. Cancer Res. DOI: 10.1158/1078-0432.CCR-18-0968.

42. Casale, E., M. Fasolini, G. Papeo, H. Posteri, D. Borghi, A. Busel, F. Caprera, M. Ciomei, A. Cirla, E. Corti, M. DAnello, M. Fasolini, E. Felder, B. Forte, A. Galvani, A. Isacchi, A. Khvat, M. Krasavin, R. Lupi, P. Orsini, R. Perego, E. Pesenti, D. Pezzetta, S. Rainoldi, F. RiccardiSirtori, A. Scolaro, F. Sola, F. Zuccotto, D. Donati, and A. Montagnoli, 2015. Structure of human PARP1 catalytic domain bound to an isoindolinone inhibitor, DOI: 10.2210/pdb4ZZZ/pdb.

43. Bossak, K., W. Goch, K. Piątek, T. Frączyk, J. Poznański, A. Bonna, C. Keil, A. Hartwig, and W. Bal, 2015. Unusual Zn(II) affinities of zinc fingers of poly(ADP-ribose) polymerase 1 (PARP-1) nuclear protein. Chem. Res. Toxicol. 28:191–201, DOI: 10.1021/tx500320f.

44. Haince, J.-F., D. McDonald, A. Rodrigue, U. Déry, J.-Y. Masson, M. J. Hendzel, and G. G. Poirier, 2008. PARP1-dependent kinetics of recruitment of MRE11 and NBS1 proteins to multiple DNA damage sites*. J. Biol. Chem. 283:1197–1208, DOI: 10.1074/jbc.M706734200.

45. Langelier, M.-F., J. L. Planck, S. Roy, and J. M. Pascal, 2011. Crystal structures of poly(ADP-ribose) polymerase-1 (PARP-1) zinc fingers bound to DNA: Structural and functional insights into DNA-dependent PARP-1 activity*,. J. Biol. Chem. 286:10690–10701, DOI: 10.1074/jbc.M110.202507.

46. Alemasova, E. E., and O. I. Lavrik, 2019. Poly(ADP-ribosyl)ation by PARP1: reaction mechanism and regulatory proteins. Nucleic Acids Res. 47:3811–3827, DOI: 10.1093/nar/gkz120.

47. Kim, M. Y., T. Zhang, and W. L. Kraus, 2005. Poly(ADP-ribosyl)ation by PARP-1: ‘PAR-laying’ NAD+ into a nuclear signal. Genes Dev. 19:1951–1967, DOI: 10.1101/gad.1331805.

48. Dawicki-McKenna, J., M.-F. Langelier, J. DeNizio, A. Riccio, C. Cao, K. Karch, M. McCauley, J. Steffen, B. Black, and J. Pascal, 2015. PARP-1 activation requires local unfolding of an autoinhibitory domain. Mol. Cell 60:755–768, DOI: 10.1016/j.molcel.2015.10.013.

49. Ogden, T. E. H., J.-C. Yang, M. Schimpl, L. E. Easton, E. Underwood, P. Rawlins, M. McCauley, M.-F. Langelier, J. Pascal, K. Embrey, and D. Neuhaus, 2021. Dynamics of the HD regulatory subdomain of PARP-1; substrate access and allostery in PARP activation and inhibition. Nucleic Acids Res. 49:2266–2288, DOI: 10.1093/nar/gkab020.

50. Singh, N., 1991. Enhanced poly ADP-ribosylation in human leukemia lymphocytes and ovarian cancers. Cancer Lett. 58:131–135, DOI: 10.1016/0304-3835(91)90035-G.

51. Nomura, F., M. Yaguchi, A. Togawa, M. Miyazaki, K. Isobe, M. Miyake, M. Noda, and T. Nakai, 2000. Enhancement of poly-adenosine diphosphate-ribosylation in human hepatocellular carcinoma. J. Gastroenterol. Hepatol. 15:529–535, DOI: 10.1046/j.1440-1746.2000.02193.x.

52. Yalcintepe, L., L. Turker-Sener, A. Sener, G. Yetkin, D. Tiryaki, and E. Bermek, 2005. Changes in NAD/ADP-ribose metabolism in rectal cancer. Braz. J. Med. Biol. Res. 38:361–365, DOI: 10.1590/S0100-879X2005000300006.

53. Ossovskaya, V., I. C. Koo, E. P. Kaldjian, C. Alvares, and B. M. Sherman, 2010. Upregulation of poly (ADP-ribose) polymerase-1 (PARP1) in triple-negative breast cancer and other primary human tumor types. Genes Cancer 1:812–821, DOI: 10.1177/1947601910383418.

54. Schiewer, M. J., J. F. Goodwin, S. Han, J. C. Brenner, M. A. Augello, J. L. Dean, F. Liu, J. L. Planck, P. Ravindranathan, A. M. Chinnaiyan, P. McCue, L. G. Gomella, G. V. Raj, A. P. Dicker, J. R. Brody, J. M. Pascal, M. M. Centenera, L. M. Butler, W. D. Tilley, F. Y. Feng, and K. E. Knudsen, 2012. Dual roles of PARP-1 promote cancer growth and progression. Cancer Discovery 2:1134–1149, DOI: 10.1158/2159-8290.CD-12-0120.

55. Bi, F.-F., D. Li, and Q. Yang, 2013. Hypomethylation of ETS transcription factor binding sites and upregulation of PARP1 expression in endometrial cancer. BioMed Res. Int. 2013:946268, DOI: 10.1155/2013/946268.

56. Green, A. R., D. Caracappa, A. A. Benhasouna, A. Alshareeda, C. C. Nolan, R. D. Macmillan, S. Madhusudan, I. O. Ellis, and E. A. Rakha, 2015. Biological and clinical significance of PARP1 protein expression in breast cancer. Breast Cancer Res. Treat. 149:353–362, DOI: 10.1007/s10549-014-3230-1.

57. Virág, L., A. L. Salzman, and C. Szabó, 1998. Poly(ADP-Ribose) synthetase activation mediates mitochondrial injury during oxidant-induced cell death. J. Immunol. 161:3753–3759.

58. Ha, H. C., and S. H. Snyder, 1999. Poly(ADP-ribose) polymerase is a mediator of necrotic cell death by ATP depletion. Proc. Natl. Acad. Sci. U. S. A. 96:13978–13982, DOI: 10.1073/pnas.96.24.13978.

59. Berger, N. A., V. C. Besson, A. H. Boulares, A. Bürkle, A. Chiarugi, R. S. Clark, N. J. Curtin, S. Cuzzocrea, T. M. Dawson, V. L. Dawson, G. Haskó, L. Liaudet, F. Moroni, P. Pacher, P. Radermacher, A. L. Salzman, S. H. Snyder, F. G. Soriano, R. P. Strosznajder, B. Sümegi, R. A. Swanson, and C. Szabo, 2018. Opportunities for the repurposing of PARP inhibitors for the therapy of non-oncological diseases. Br. J. Pharmacol. 175:192–222, DOI: 10.1111/bph.13748.

60. Wang, X.-G., Z.-Q. Wang, W.-M. Tong, and Y. Shen, 2007. PARP1 Val762Ala polymorphism reduces enzymatic activity. Biochem. Biophys. Res. Commun. 354:122–126, DOI: 10.1016/j.bbrc.2006.12.162.

61. Ma, X.-B., X.-J. Wang, M. Wang, Z.-M. Dai, T.-B. Jin, X.-H. Liu, H.-F. Kang, S. Lin, P. Xu, and Z.-J. Dai, 2016. Impact of the PARP1 rs1136410 and rs3219145 polymorphisms on susceptibility and clinicopathologic features of breast cancer in a Chinese population. Transl. Cancer Res. 5, DOI: 10.21037/tcr.2016.09.01.

62. Wang, X.-b., N.-h. Cui, S. Zhang, S.-r. Guo, Z.-j. Liu, and L. Ming, 2017. PARP-1 variant rs1136410 confers protection against coronary artery disease in a Chinese Han population: A two-stage case-control study involving 5643 subjects. Front. Physiol. 8:916, DOI: 10.3389/fphys.2017.00916.

63. Cui, N.-H., C. Qiao, X.-K. Chang, and L. Wei, 2017. Associations of PARP-1 variant rs1136410 with PARP activities, oxidative DNA damage, and the risk of age-related cataract in a Chinese Han population: A two-stage case-control analysis. Gene 600:70–76, DOI: 10.1016/j.gene.2016.11.019.

64. Li, H., Y. Zha, F. Du, J. Liu, X. Li, and X. Zhao, 2020. Contributions of PARP-1 rs1136410 C>T polymorphism to the development of cancer. J. Cell. Mol. Med. 24:14639–14644, DOI: 10.1111/jcmm.16027.

65. Xin, Y., L. Yang, M. Su, X. Cheng, L. Zhu, and J. Liu, 2021. PARP1 rs1136410 Val762Ala contributes to an increased risk of overall cancer in the East Asian population: a metaanalysis. J. Int. Med. Res. 49:0300060521992956, DOI: 10.1177/0300060521992956.

66. El-Gebali, S., J. Mistry, A. Bateman, S. R. Eddy, A. Luciani, S. C. Potter, M. Qureshi, L. J. Richardson, G. A. Salazar, A. Smart, E. L. L. Sonnhammer, L. Hirsh, L. Paladin, D. Piovesan, S. C. E. Tosatto, and R. D. Finn, 2018. The Pfam protein families database in 2019. Nucleic Acids Res. 47, DOI: 10.1093/nar/gky995.

67. Figliuzzi, M., P. Barrat-Charlaix, and M. Weigt, 2018. How pairwise coevolutionary models capture the collective residue variability in proteins? Mol. Biol. Evol. 35:1018–1027, DOI: 10.1093/molbev/msy007.

68. Trinquier, J., G. Uguzzoni, A. Pagnani, F. Zamponi, and M. Weigt, 2021. Efficient generative modeling of protein sequences using simple autoregressive models. Nat. Commun. 12, DOI: 10.1038/s41467-021-25756-4.

69. Cheng, R. R., O. Nordesjö, R. L. Hayes, H. Levine, S. C. Flores, J. N. Onuchic, and F. Morcos, 2016. Connecting the sequence-space of bacterial signaling proteins to phenotypes using coevolutionary landscapes. Mol. Biol. Evol. 33:3054–3064, DOI: 10.1093/molbev/msw188.

70. Figliuzzi, M., H. Jacquier, A. Schug, O. Tenaillon, and M. Weigt, 2015. Coevolutionary landscape inference and the context-dependence of mutations in Beta-Lactamase TEM-1. Mol. Biol. Evol. 33:268–280, DOI: 10.1093/molbev/msv211.

71. Zhou, Q., N. Kunder, J. A. D. la Paz, A. E. Lasley, V. D. Bhat, F. Morcos, and Z. T. Campbell, 2018. Global pairwise RNA interaction landscapes reveal core features of protein recognition. Nat. Commun. 9, DOI: 10.1038/s41467-018-04729-0.

72. Jiang, X.-L., R. P. Dimas, C. T. Y. Chan, and F. Morcos, 2021. morcoslab/coevolution-compatibility: Companion to “Coevolutionary methods enable robust design of modular repressors by reestablishing intra-protein interactions”. DOI: 10.5281/zenodo.5262799, DOI: 10.5281/zenodo.5262799.

73. Papeo, G., H. Posteri, D. Borghi, A. A. Busel, F. Caprera, E. Casale, M. Ciomei, A. Cirla, E. Corti, M. D’Anello, M. Fasolini, B. Forte, A. Galvani, A. Isacchi, A. Khvat, M. Y. Krasavin, R. Lupi, P. Orsini, R. Perego, E. Pesenti, D. Pezzetta, S. Rainoldi, F. Riccardi-Sirtori, A. Scolaro, F. Sola, F. Zuccotto, E. R. Felder, D. Donati, and A. Montagnoli, 2015. Discovery of 2-[1-(4,4-Difluorocyclohexyl)piperidin-4-yl]-6-fluoro-3-oxo-2,3-dihydro-1H-isoindole-4-carboxamide (NMS-P118): A potent, orally available, and highly selective PARP-1 inhibitor for cancer therapy. J. Med. Chem. 58:6875–6898, DOI: 10.1021/acs.jmedchem.5b00680.

74. Cao, R., Y. Wang, J. Zhou, N. Huang, and B. Xu, 2016. Structure of human PARP1 catalytic domain bound to a benzoimidazole inhibitor, DOI: 10.2210/pdb5WS1/pdb.

75. Schafmeister, C. E. A. F., W. S. Ross, and V. Romanovski, 1995. LEaP.

76. Case, D., I. Ben-Shalom, S. Brozell, D. Cerutti, T. Cheatham III, V. Cruzeiro, T. Darden, R. Duke, D. Ghoreishi, M. Gilson, H. Gohlke, A. Goetz, D. Greene, R. Harris, N. Homeyer, Y. Huang, S. Izadi, A. Kovalenko, T. Kurtzman, T. Lee, S. LeGrand, P. Li, C. Lin, J. Liu, T. Luchko, R. Luo, D. Mermelstein, K. Merz, Y. Miao, G. Monard, C. Nguyen, H. Nguyen, I. Omelyan, A. Onufriev, F. Pan, R. Qi, D. R. Roe, A. Roitberg, C. Sagui, S. Schott-Verdugo, J. Shen, C. Simmerling, J. Smith, R. Salomon-Ferrer, J. Swails, R. Walker, J. Wang, H. Wei, R. Wolf, X. Wu, L. Xiao, D. York, and P. Kollman, 2018. AMBER 2018.

77. Šali, A., and T. L. Blundell, 1993. Comparative protein modelling by satisfaction of spatial restraints. J. Mol. Biol. 234:779 – 815, DOI: 10.1006/jmbi.1993.1626.

78. Fiser, A., R. K. G. Do, and A. Šali, 2000. Modeling of loops in protein structures. Protein Sci. 9:1753–1773, DOI: 10.1110/ps.9.9.1753.

79. Pettersen, E. F., T. D. Goddard, C. C. Huang, G. S. Couch, D. M. Greenblatt, E. C. Meng, and T. E. Ferrin, 2004. UCSF Chimera—A visualization system for exploratory research and analysis. J. Comput. Chem. 25:1605–1612, DOI: 10.1002/jcc.20084.

80. Dunbrack, R. L., 2002. Rotamer libraries in the 21st century. Curr. Opin. Struct. Biol. 12:431–440, DOI: 10.1016/S0959-440X(02)00344-5.

81. Humphrey, W., A. Dalke, and K. Schulten, 1996. VMD: Visual molecular dynamics. J. Mol. Graphics 14:33 – 38, DOI: 10.1016/0263-7855(96)00018-5.

82. Jorgensen, W. L., J. Chandrasekhar, J. D. Madura, R. W. Impey, and M. L. Klein, 1983. Comparison of simple potential functions for simulating liquid water. J. Chem. Phys. 79:926–935, DOI: 10.1063/1.445869.

83. Søndergaard, C. R., M. H. M. Olsson, M. Rostkowski, and J. H. Jensen, 2011. Improved treatment of ligands and coupling effects in empirical calculation and rationalization of pKa values. J. Chem. Theory Comput. 7:2284–2295, DOI: 10.1021/ct200133y.

84. Olsson, M. H. M., C. R. Søndergaard, M. Rostkowski, and J. H. Jensen, 2011. PROPKA3: Consistent treatment of internal and surface residues in empirical pKa predictions. J. Chem. Theory Comput. 7:525–537, DOI: 10.1021/ct100578z.

85. Jurrus, E., D. Engel, K. Star, K. Monson, J. Brandi, L. E. Felberg, D. H. Brookes, L. Wilson, J. Chen, K. Liles, M. Chun, P. Li, D. W. Gohara, T. Dolinsky, R. Konecny, D. R. Koes, J. E. Nielsen, T. Head-Gordon, W. Geng, R. Krasny, G.-W. Wei, M. J. Holst, J. A. McCammon, and N. A. Baker, 2018. Improvements to the APBS biomolecular solvation software suite. Protein Sci. 27:112–128, DOI: 10.1002/pro.3280.

86. Maier, J. A., C. Martinez, K. Kasavajhala, L. Wickstrom, K. E. Hauser, and C. Simmerling, 2015. ff14SB: Improving the accuracy of protein side chain and backbone parameters from ff99SB. J. Chem. Theory Comput. 11:3696–3713, DOI: 10.1021/acs.jctc.5b00255.

87. Essmann, U., L. Perera, M. L. Berkowitz, T. Darden, H. Lee, and L. G. Pedersen, 1995. A smooth particle mesh Ewald method. J. Chem. Phys. 103:8577–8593, DOI: 10.1063/1.470117.

88. Salomon-Ferrer, R., A. W. Götz, D. Poole, S. Le Grand, and R. C. Walker, 2013. Routine microsecond molecular dynamics simulations with AMBER on GPUs. 2. Explicit solvent particle mesh Ewald. J. Chem. Theory Comput. 9:3878–3888, DOI: 10.1021/ct400314y.

89. Loncharich, R. J., B. R. Brooks, and R. W. Pastor, 1992. Langevin dynamics of peptides: The frictional dependence of isomerization rates of N-acetylalanyl-N’-methylamide. Biopolymers 32:523–535, DOI: 10.1002/bip.360320508.

90. Roe, D. R., and T. E. Cheatham, 2013. PTRAJ and CPPTRAJ: Software for processing and analysis of molecular dynamics trajectory data. J. Chem. Theory Comput. 9:3084– 3095, DOI: 10.1021/ct400341p.

91. Bakan, A., I. Bahar, and L. M. Meireles, 2011. ProDy: Protein dynamics inferred from theory and experiments. Bioinformatics 27:1575–1577, DOI: 10.1093/bioinformatics/btr168.

92. Williams, T., and C. Kelley, 2019. Gnuplot 5.2: An interactive plotting program. http://gnuplot.sourceforge.net/.

93. Hunter, J. D., 2007. Matplotlib: A 2D graphics environment. Computing in Science & Engineering 9:90–95, DOI: 10.1109/MCSE.2007.55.

94. Oliphant, T. E., 2006. A guide to NumPy, volume 1. Trelgol Publishing USA.

95. Van Der Walt, S., S. C. Colbert, and G. Varoquaux, 2011. The NumPy array: A structure for efficient numerical computation. Computing in Science & Engineering 13:22.

96. Seabold, S., and J. Perktold, 2010. statsmodels: Econometric and statistical modeling with python. In 9th Python in Science Conference.

97. Leddin, E., and G. A. Cisneros, 2020. CisnerosResearch/AMBER-EDA: First Release. DOI: 10.5281/zenodo.4469902, DOI: 10.5281/zenodo.4469902.

98. R Core Team, 2018. R: A language and environment for statistical computing. R Foundation for Statistical Computing, Vienna, Austria. https://www.R-project.org/.

99. Dowle, M., and A. Srinivasan, 2018. data.table: Extension of ‘data.frame’. https://CRAN.R-project.org/package=data.table, r package version 1.11.8.

100. Plate, T., and R. Heiberger, 2016. abind: Combine multidimensional arrays. https://CRAN.R-project.org/package=abind, r package version 1.4-5.

101. Wickham, H., 2017. tidyverse: Easily install and load the ‘Tidyverse’. https://CRAN.R-project.org/package=tidyverse, r package version 1.2.1.

