## Supporting Material for "Computational Compensatory Mutation Discovery Approach: Predicting a PARP1 Variant Rescue Mutation"

Krithika Ravishankar[1, ‡], Xianli Jiang[2,3,‡], Emmett M. Leddin[1], Faruck Morcos[2,4,5,*], and G. Andrés Cisneros[1,6,7,*]

[1]Department of Chemistry, University of North Texas, Denton, TX 76201
[2]Department of Biological Sciences, The University of Texas at Dallas, Richardson, TX 75080
[3]Department of Bioinformatics and Computational Biology, The University of Texas MD Anderson Cancer Center
[4]Department of Bioengineering, The University of Texas at Dallas, Richardson, TX 75080
[5]Center for Systems Biology, The University of Texas at Dallas, Richardson, TX 75080
[6]Department of Physics, The University of Texas at Dallas, Richardson, TX 75080
[7]Department of Chemistry, The University of Texas at Dallas, Richardson, TX 75080
[‡]Authors have equivalent contributions
[*]

April 18, 2022

### Table of Contents


**1. Coevolutionary Analysis** **5**

**2. RMSD and RMSF** **6**
2.1. RMSD Per System 6
2.2. RMSF Per System 9
2.3. RMSD and RMSF for a Representative Trajectory of Each System 12

**3. Correlation Matrices** **14**
3.1. V762A 14
3.2. V762A-from-WT 14
3.3. A755E 15
3.4. A755L 15
3.5. A755E/V762A 16
3.6. A755L/V762A 16

**4. Energy Decomposition Analysis (EDA)** **17**
4.1. WT 17
4.2. V762A 18
4.3. V762A-from-WT 19
4.4. A755E 20
4.5. A755L 21
4.6. A755E/V762A 22
4.7. A755L/V762A 24
4.8. Difference EDA 26

**5. Normal Mode Analysis** **28**

### List of Figures


S1. Double Mutation landscape calculated from coevolutionary information based energy for PARP1 SNP V762A. Each data point represents the H(S) score for a double mutant that contains V762A. 5
S2. Root mean square deviations (RMSD) for the WT and V762A variant systems. Bézier line smoothing has been applied. 6
S3. Root mean square deviations (RMSD) for the A755E and A755L single- and double- variant systems (with V762A). Bézier line smoothing has been applied. 7
S4. Root mean square deviations (RMSD) for the WT system with TIP3P water extending 12 Å from the protein surface. Raw data are shown in pink, with line smoothing applied in white. 8
S5. Root mean square fluctuations (RMSF) for the WT and V762A variant systems. 9
S6. Root mean square fluctuations (RMSF) for the A755E and A755L single- and double- variant systems (with V762A). 10
S7. Root mean square fluctuations (RMSF) for the WT system with TIP3P water extending 12 Å from the protein surface. 11
S8. RMSD, RMSF, and Difference RMSF (Variant – WT) for a representative trajectory of each system plotted together. 12
S9. Correlation matrices of V762A. Areas of correlation are blue, areas with no correlation are yellow, and areas with anti-correlation are red. 14
S10. Correlation matrices of V762A-from-WT. Areas of correlation are blue, areas with no correlation are yellow, and areas with anti-correlation are red. 14
S11. Correlation matrices of A755E. Areas of correlation are blue, areas with no correlation are yellow, and areas with anti-correlation are red. 15
S12. Correlation matrices of A755L. Areas of correlation are blue, areas with no correlation are yellow, and areas with anti-correlation are red. 15
S13. Correlation matrices of A755E/V762A. Areas of correlation are blue, areas with no correlation are yellow, and areas with anti-correlation are red. 16
S14. Correlation matrices of A755L/V762A. Areas of correlation are blue, areas with no correlation are yellow, and areas with anti-correlation are red. 16
S15. Total interaction energy (Coulomb and van der Waals) with respect to residue 762 in WT. 17
S16. Total interaction energy (Coulomb and van der Waals) with respect to residue 762 in V762A. 18
S17. Total interaction energy (Coulomb and van der Waals) with respect to residue 762 in V762A-from-WT. 19
S18. Total interaction energy (Coulomb and van der Waals) with respect to residue 762 in A755E. 20
S19. Total interaction energy (Coulomb and van der Waals) with respect to residue 762 in A755L. 21
S20. Total interaction energy (Coulomb and van der Waals) with respect to residue 755 (highlighted in gray) in A755E/V762A. 22
S21. Total interaction energy (Coulomb and van der Waals) with respect to residue 762 (highlighted in gray) in A755E/V762A. 23

S22. Total interaction energy (Coulomb and van der Waals) with respect to residue 755 (highlighted in gray) in A755L/V762A. 24
S23. Total interaction energy (Coulomb and van der Waals) with respect to residue 762 (highlighted in gray) in A755L/V762A. 25
S24. Difference in average total interaction energy (Coulomb and van der Waals) for the system for the variant – WT. V762A and V762A-from-WT are with respect to the interactions for residue 762, and A755E and A755L are with respect to the interactions for residue 755. The variant position is highlighted gray, and average standard deviation is indicated with black error bars. 26
S25. Difference in average total interaction energy (Coulomb and van der Waals) for the system for the double variant – WT. Each are with respect to the residue specified in the subcaption. The variant positions are in highlighted gray, and average standard deviation is indicated with black error bars. 27
S26. First normal mode for a representative simulation of each system. 28
S27. First normal mode for a representative simulation of each system. 29

### 1. Coevolutionary Analysis

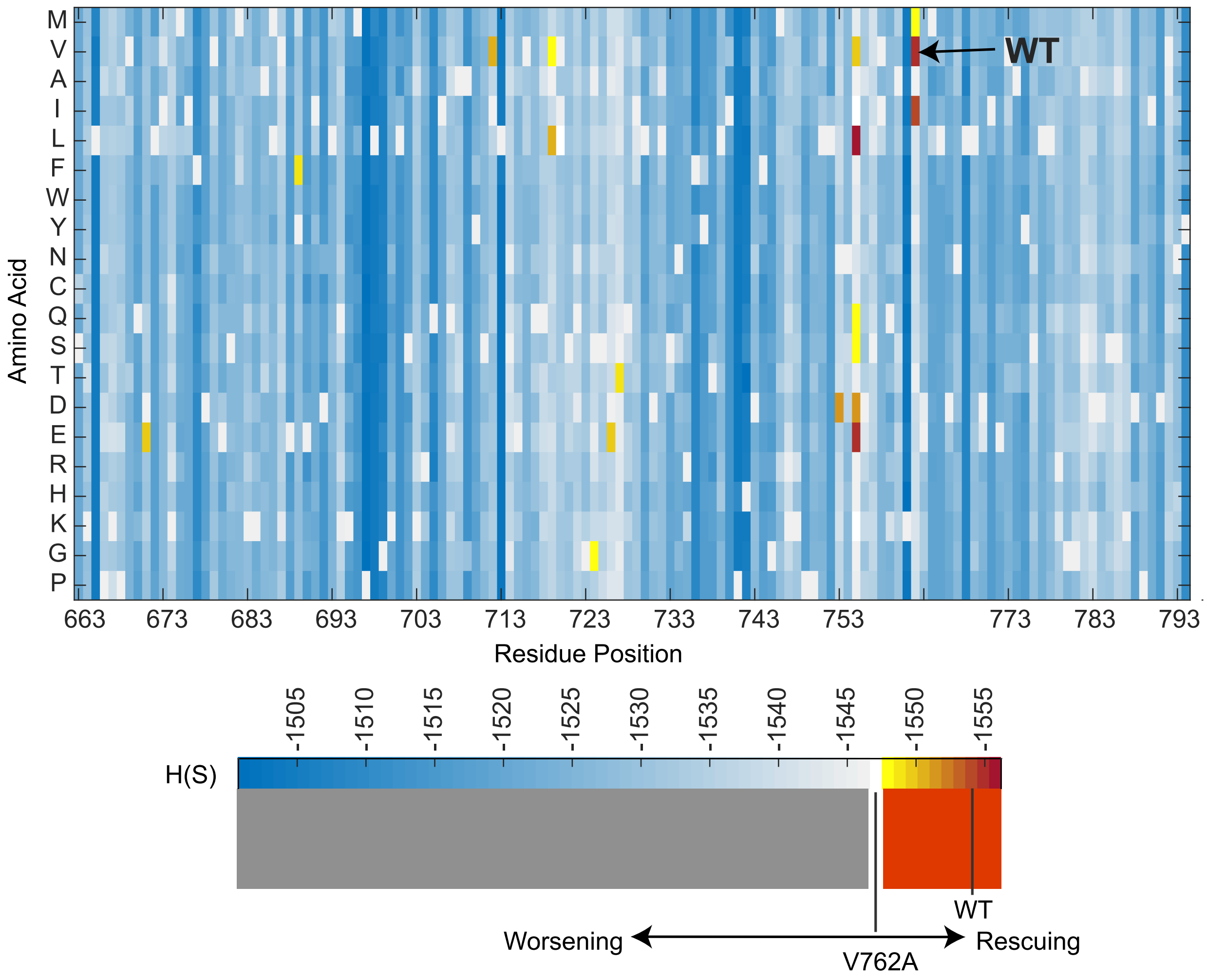


Figure S1: Double Mutation landscape calculated from coevolutionary information based energy for PARP1 SNP V762A. Each data point represents the H(S) score for a double mutant that contains V762A.

### 2. RMSD and RMSF

#### 2.1. RMSD Per System

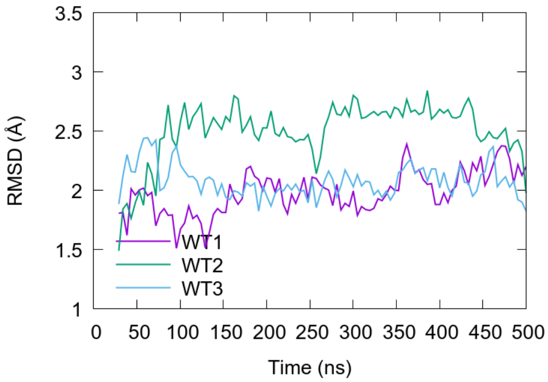


(A) WT

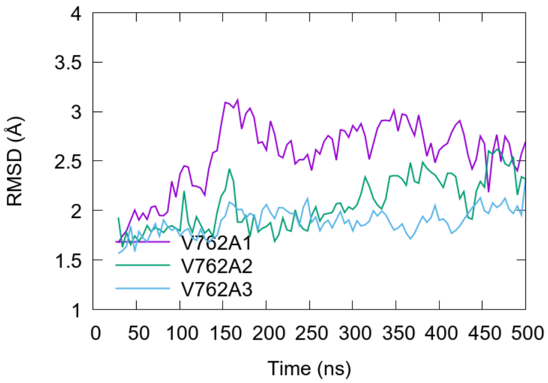


(B) V762A

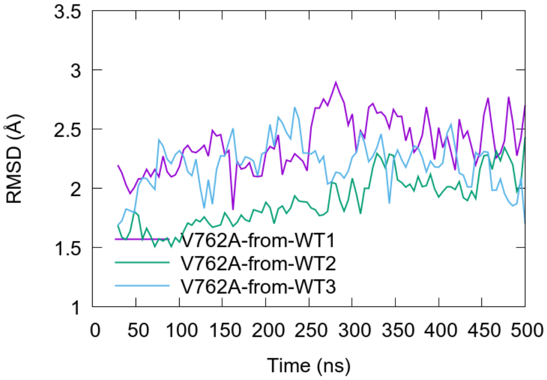


(C) V762A-from-WT

Figure S2: Root mean square deviations (RMSD) for the WT and V762A variant systems. Bézier line smoothing has been applied.

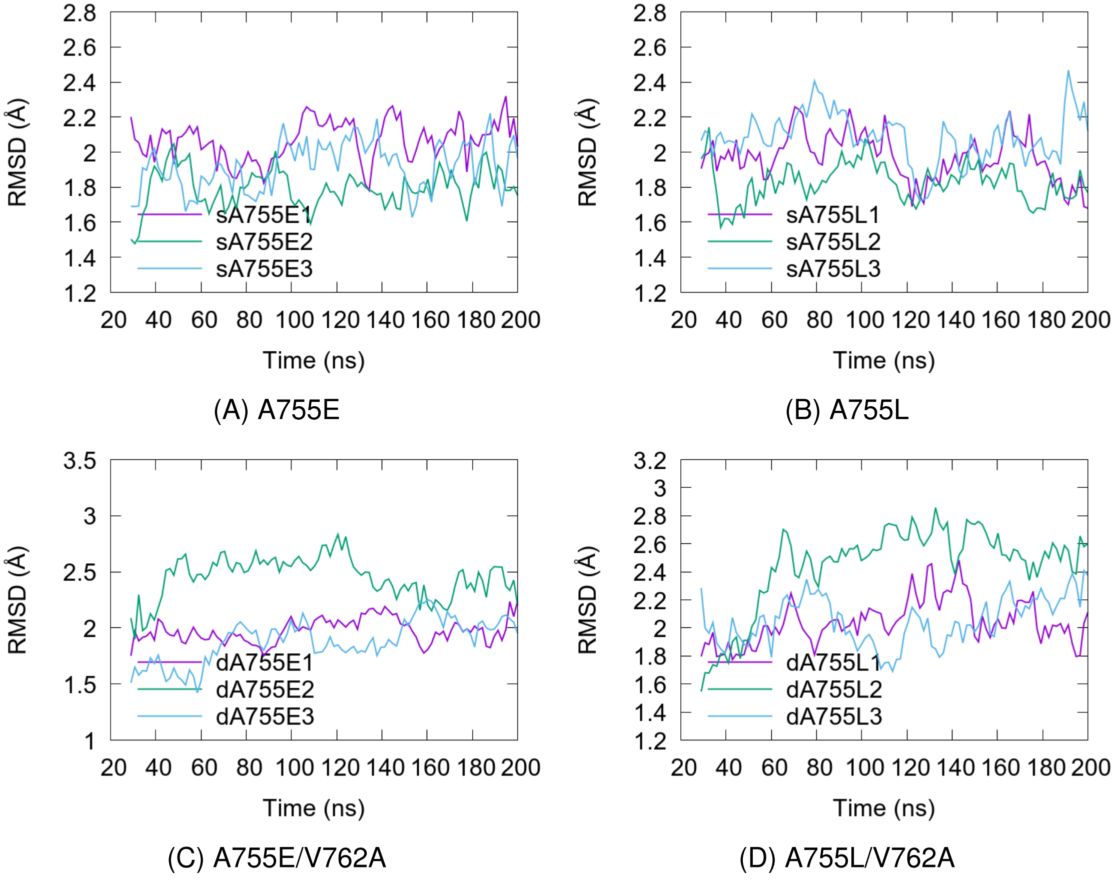


(A) A755E

(B) A755L

(C) A755E/V762A

(D) A755L/V762A

Figure S3: Root mean square deviations (RMSD) for the A755E and A755L single- and double-variant systems (with V762A). Bézier line smoothing has been applied.

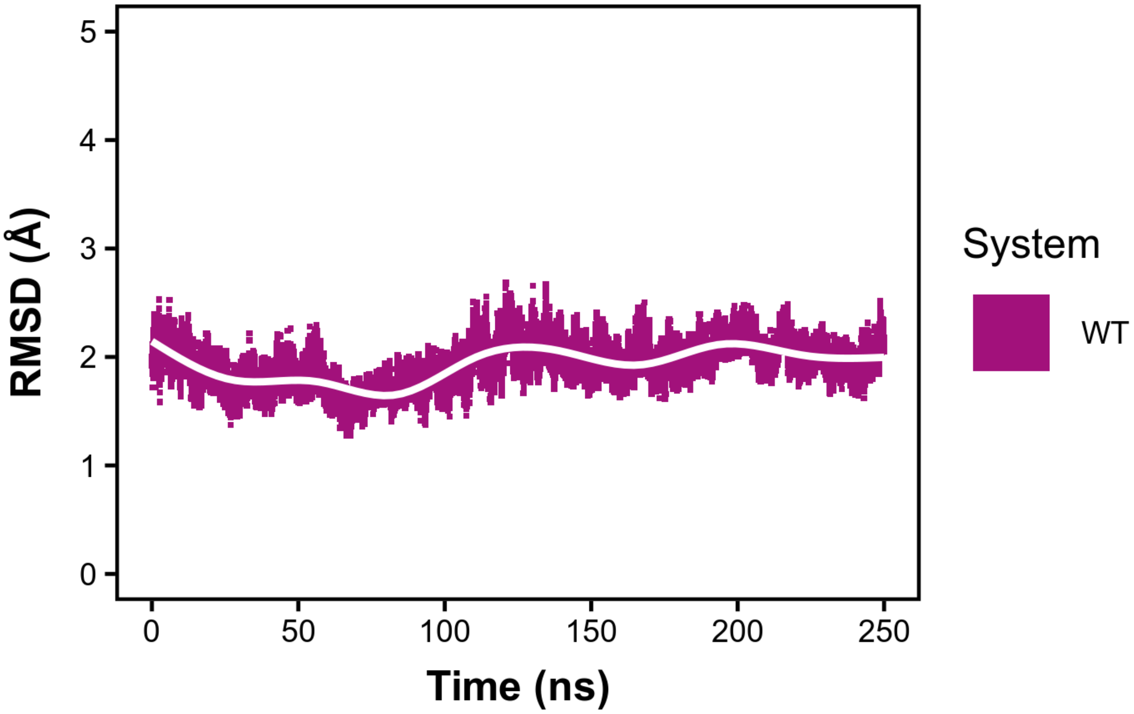


Figure S4: Root mean square deviations (RMSD) for the WT system with TIP3P water extending 12 Å from the protein surface. Raw data are shown in pink, with line smoothing applied in white.

#### 2.2. RMSF Per System

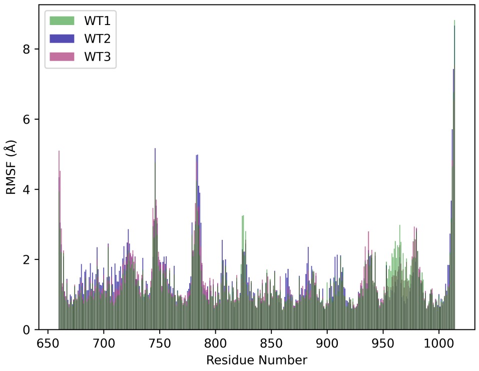


(A) WT

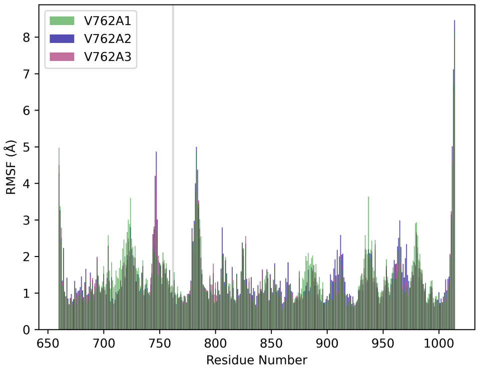


(B) V762A

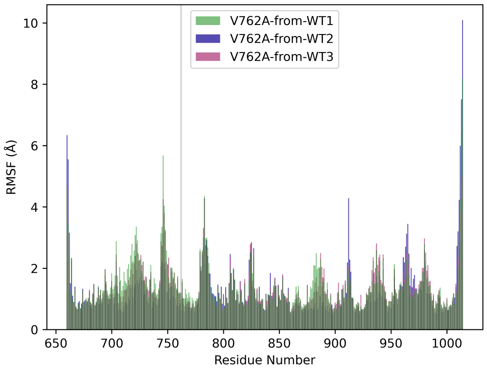


(C) V762A-from-WT

Figure S5: Root mean square fluctuations (RMSF) for the WT and V762A variant systems.

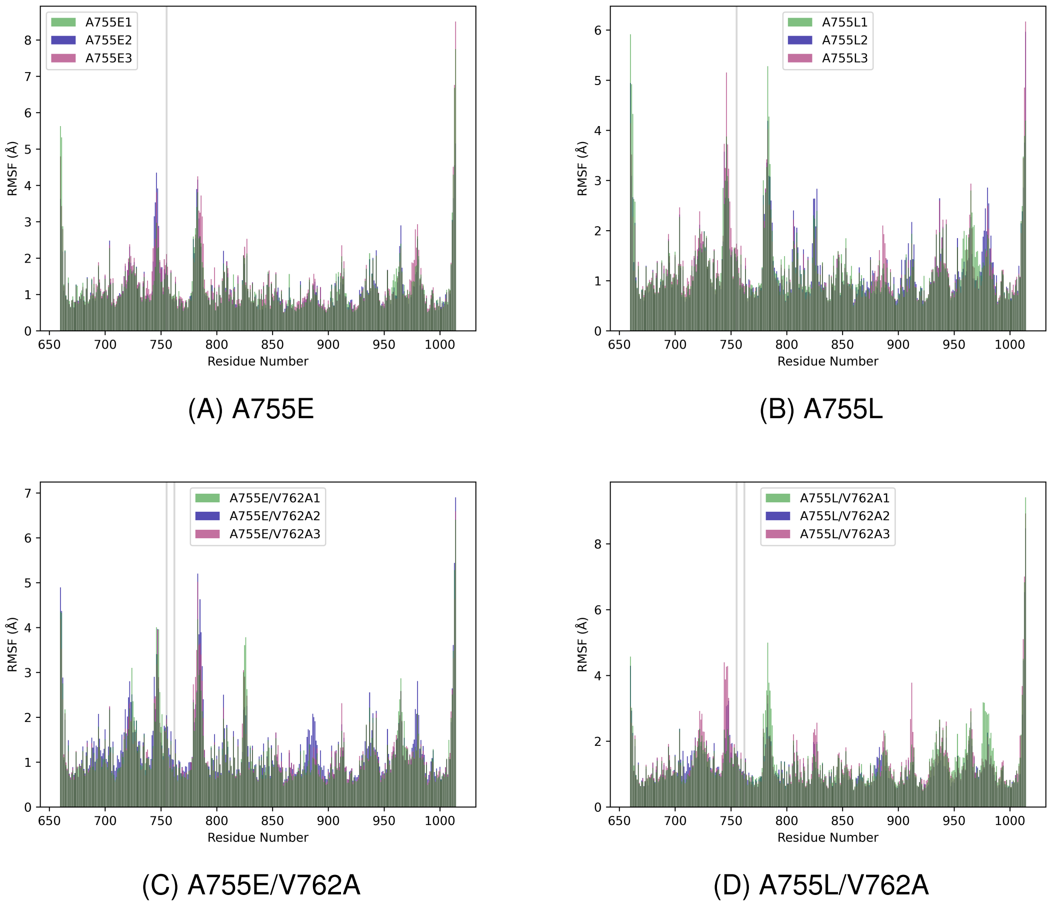


Figure S6: Root mean square fluctuations (RMSF) for the A755E and A755L single- and double- variant systems (with V762A).

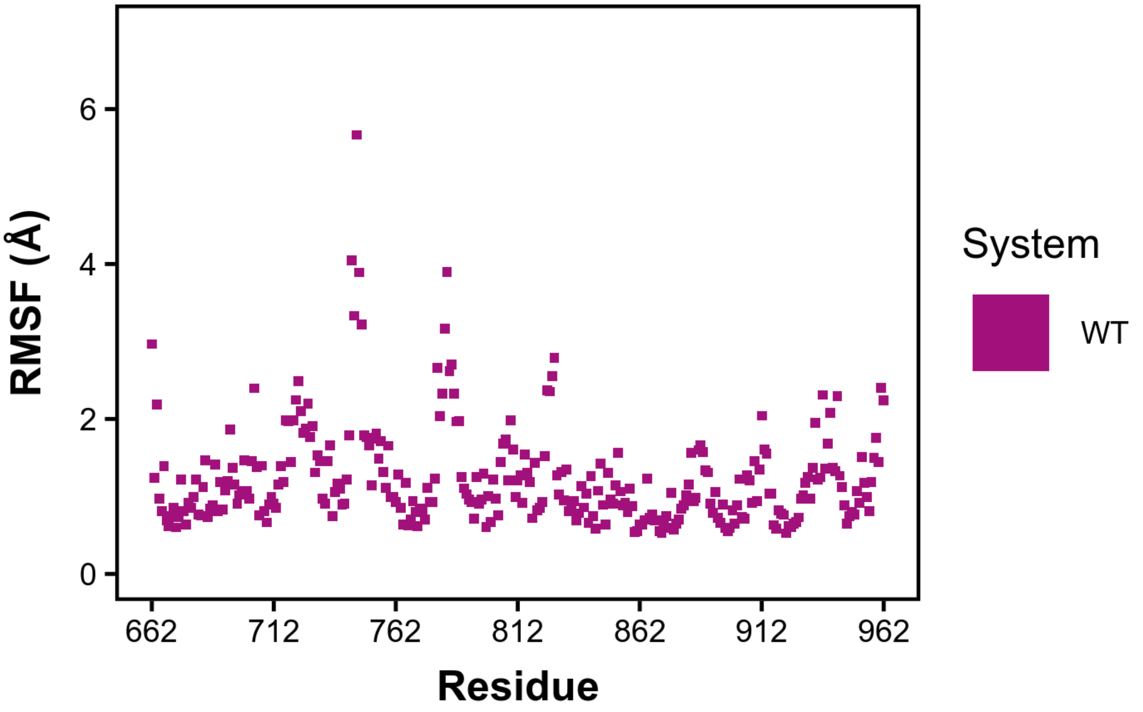


Figure S7: Root mean square fluctuations (RMSF) for the WT system with TIP3P water extending 12 Å from the protein surface.

#### 2.3. RMSD and RMSF for a Representative Trajectory of Each System

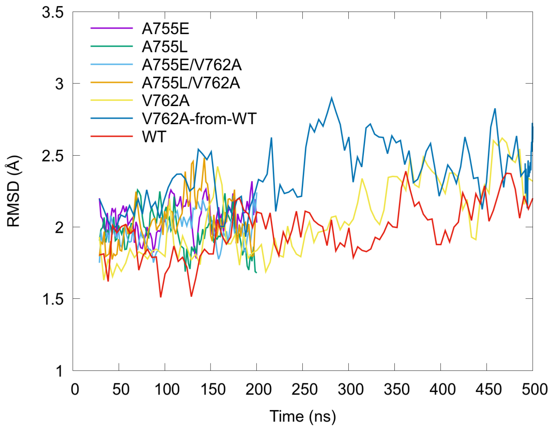


(A) RMSDs

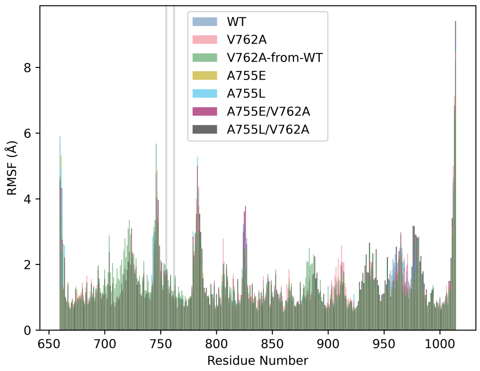


(B) RMSFs

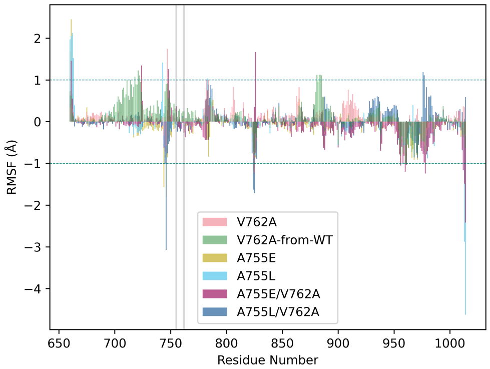


(C) Differences in RMSFs

Figure S8: RMSD, RMSF, and Difference RMSF (Variant – WT) for a representative trajectory of each system plotted together.

Table S1: Significant differences in root mean square fluctuations, variants minus WT. The helical subdomain is shaded orange (residues 710–770), and the ART subdomain is shaded red (residues 910–960).

| | V762A | V762A-from-WT | A755E | A755L | A755E/V762A | A755L/V762A |
|---|---|---|---|---|---|---|
| 660 | | | 1.69 | 1.97 | | |
| 662 | | | 2.43 | | 1.46 | |
| 663 | | | | 1.52 | | |
| 714 | | 1.00 | | | | |
| 718 | | 1.08 | | | | |
| 721 | | 1.23 | | | | |
| 722 | | 1.11 | | | | |
| 724 | 1.04 | | | | 1.35 | |
| 743 | | | | 1.42 | | |
| 744 | | | -1.57 | | | |
| 745 | | | | | | -1.01 |
| 746 | | | -1.80 | | | -3.07 |
| 747 | 1.75 | | | | | |
| 748 | | | | | 1.26 | |
| 782 | 1.01 | | | | | |
| 783 | | | | 1.03 | | |
| 824 | | | -1.17 | | | -1.63 |
| 825 | -1.23 | | -1.25 | -1.02 | | -1.71 |
| 826 | | | | | 1.67 | |
| 881 | | 1.12 | | | | |
| 883 | | 1.12 | | | | |
| 884 | | 1.12 | | | | |
| 885 | | 1.11 | | | | |
| 956 | | | | | -1.05 | |
| 960 | | -1.03 | | | -1.20 | |
| 961 | | -1.01 | | | -1.03 | |
| 976 | | | | | | 1.19 |
| 977 | | | | | | 1.11 |
| 978 | | | | | -1.24 | |
| 1012 | | | | -1.17 | -1.16 | |
| 1013 | | | | -2.88 | -1.49 | |
| 1014 | | | -1.07 | -4.63 | -2.42 | |

### 3. Correlation Matrices

## 3.1. V762A

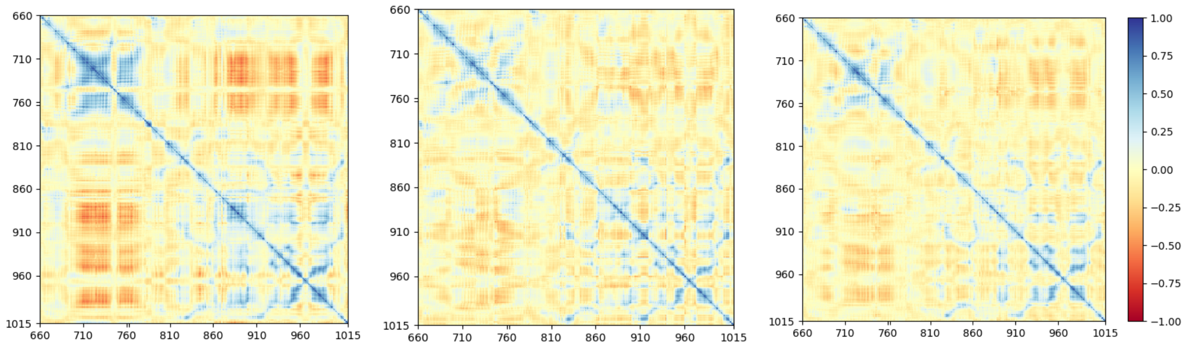


(A) V762A1 (B) V762A2 (C) V762A3

Figure S9: Correlation matrices of V762A. Areas of correlation are blue, areas with no correlation are yellow, and areas with anti-correlation are red.

#### 3.2. V762A-from-WT

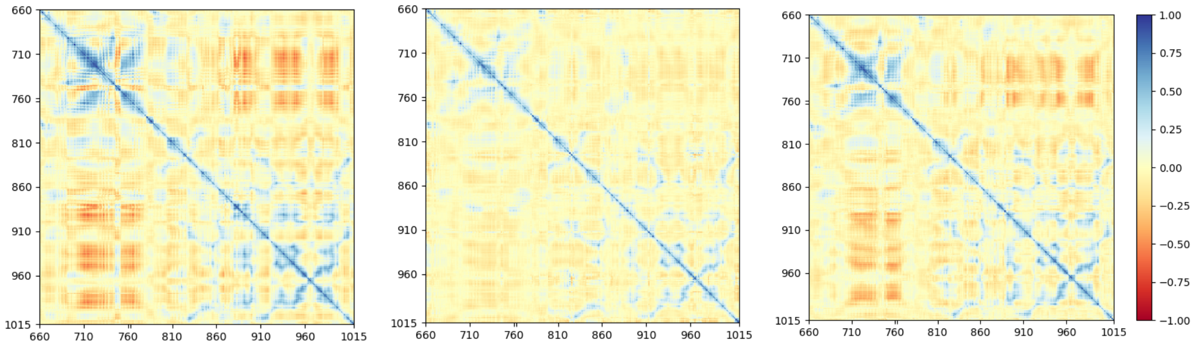


(A) V762A-from-WT1 (B) V762A-from-WT2 (C) V762A-from-WT3

Figure S10: Correlation matrices of V762A-from-WT. Areas of correlation are blue, areas with no correlation are yellow, and areas with anti-correlation are red.

## 3.3. A755E

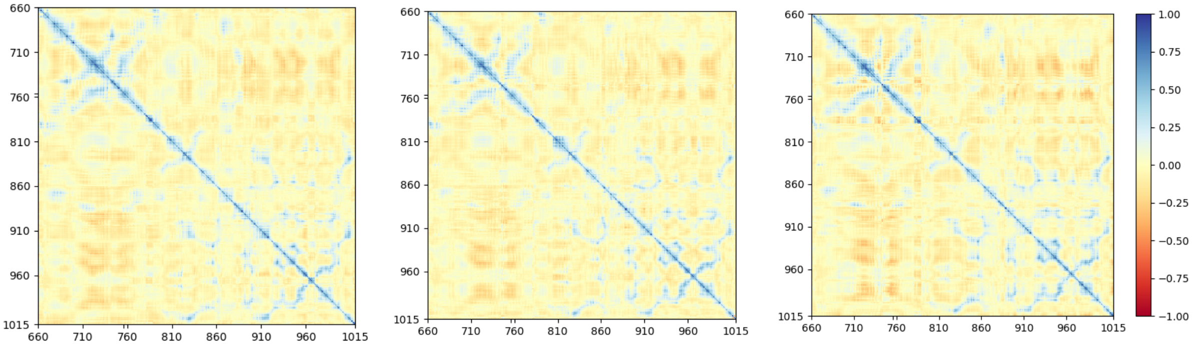

(A) A755E1 (B) A755E2 (C) A755E3

Figure S11: Correlation matrices of A755E. Areas of correlation are blue, areas with no correlation are yellow, and areas with anti-correlation are red.

## 3.4. A755L

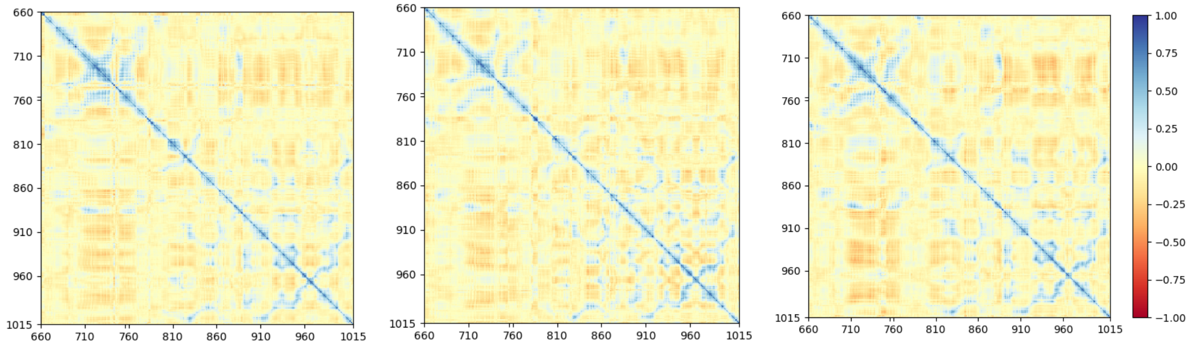

(A) A755L1 (B) A755L2 (C) A755L3

Figure S12: Correlation matrices of A755L. Areas of correlation are blue, areas with no correlation are yellow, and areas with anti-correlation are red.

## 3.5. A755E/V762A

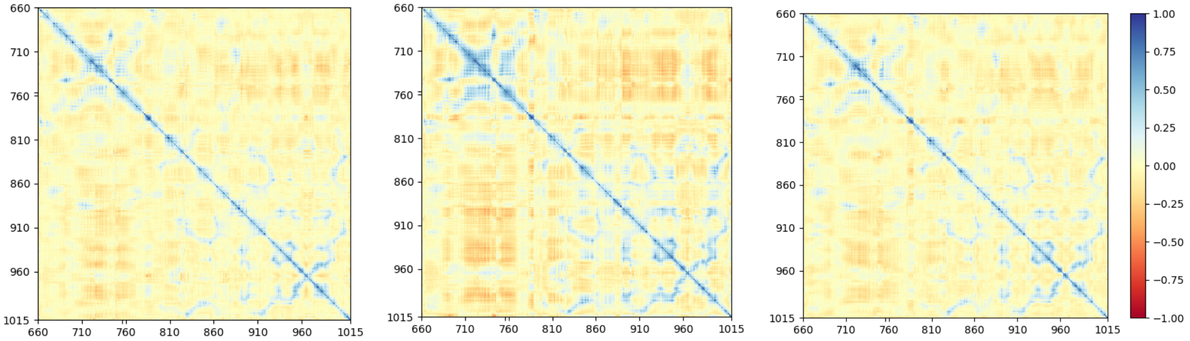


(A) A755E/V762A1

(B) A755E/V762A2

(C) A755E/V762A3

Figure S13: Correlation matrices of A755E/V762A. Areas of correlation are blue, areas with no correlation are yellow, and areas with anti-correlation are red.

## 3.6. A755L/V762A

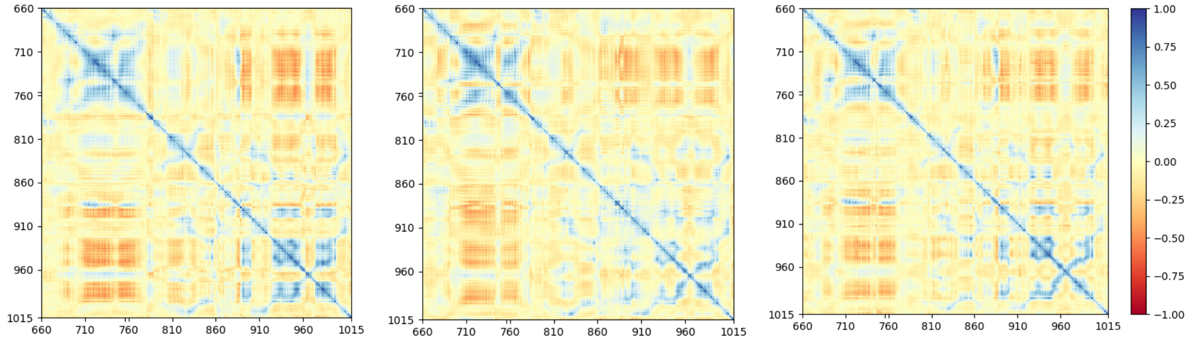


(A) A755L/V762A1

(B) A755L/V762A2

(C) A755L/V762A3

Figure S14: Correlation matrices of A755L/V762A. Areas of correlation are blue, areas with no correlation are yellow, and areas with anti-correlation are red.

### 4. Energy Decomposition Analysis (EDA)

## 4.1. WT

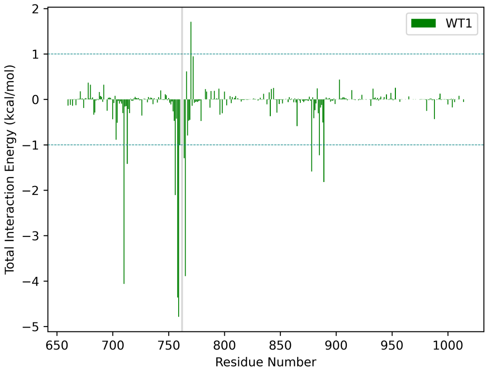


(A) WT1

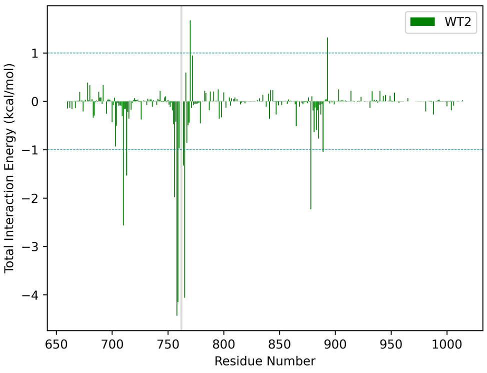


(B) WT2

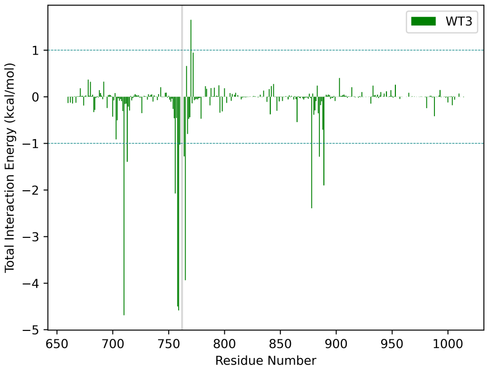


(C) WT3

Figure S15: Total interaction energy (Coulomb and van der Waals) with respect to residue 762 in WT.

## 4.2. V762A

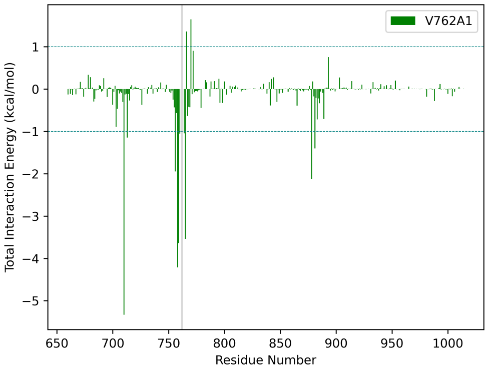


(A) V762A1

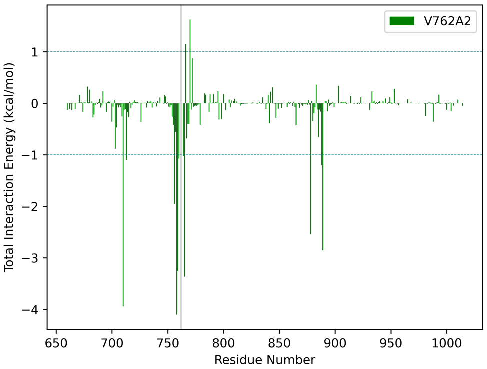


(B) V762A2

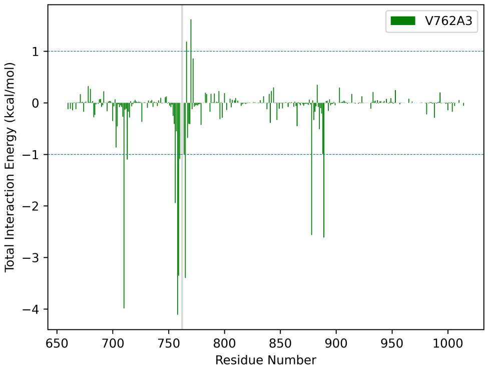


(C) V762A3

Figure S16: Total interaction energy (Coulomb and van der Waals) with respect to residue 762 in V762A.

#### 4.3. V762A-from-WT

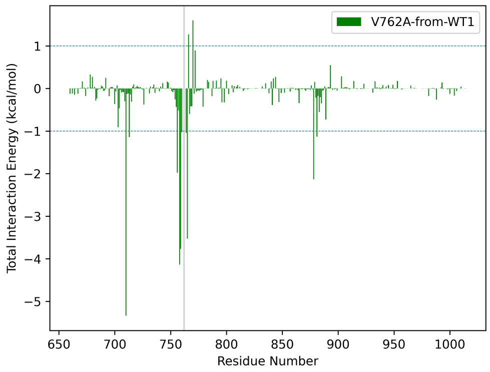


(A) V762A-from-WT1

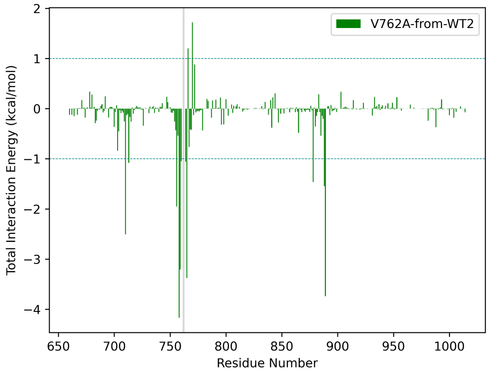


(B) V762A-from-WT2

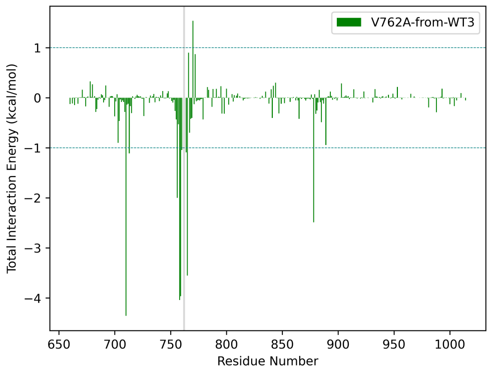


(C) V762A-from-WT3

Figure S17: Total interaction energy (Coulomb and van der Waals) with respect to residue 762 in V762A-from-WT.

## 4.4. A755E

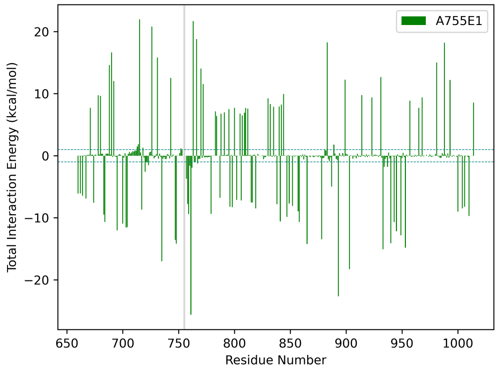


(A) A755E1

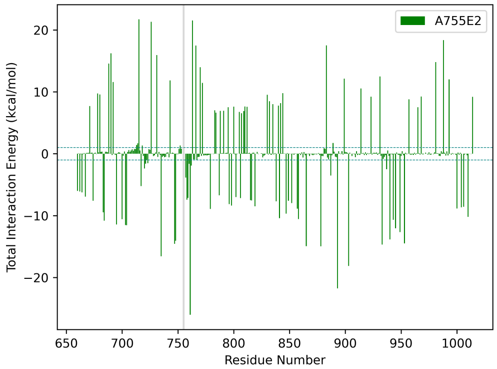


(B) A755E2

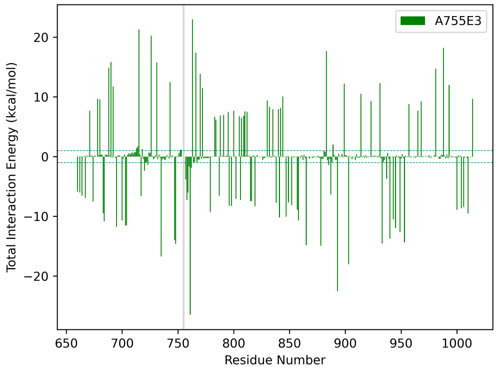


(C) A755E3

Figure S18: Total interaction energy (Coulomb and van der Waals) with respect to residue 762 in A755E.

### 4.5. A755L

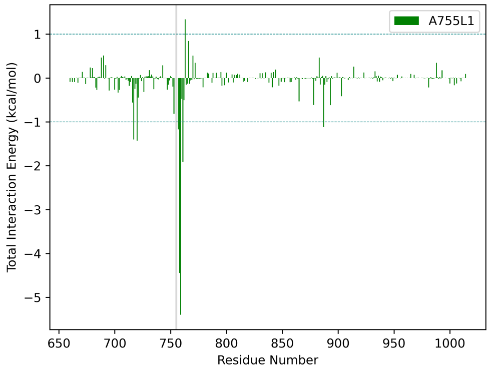


(A) A755L1

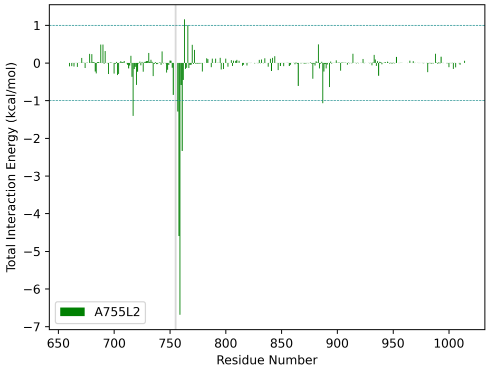


(B) A755L2

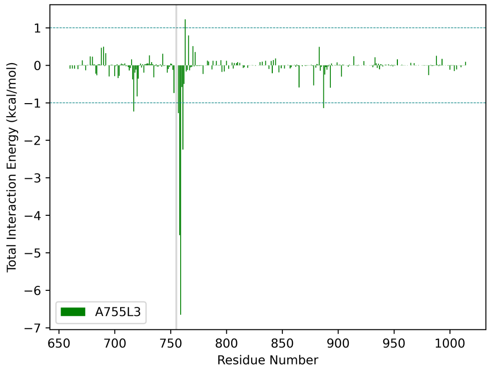


(C) A755L3

Figure S19: Total interaction energy (Coulomb and van der Waals) with respect to residue 762 in A755L.

## 4.6. A755E/V762A

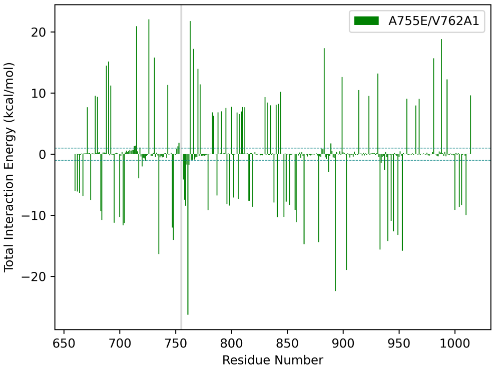


(A) A755E/V762A1

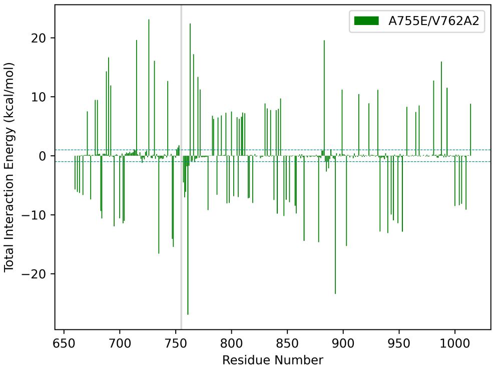


(B) A755E/V762A2

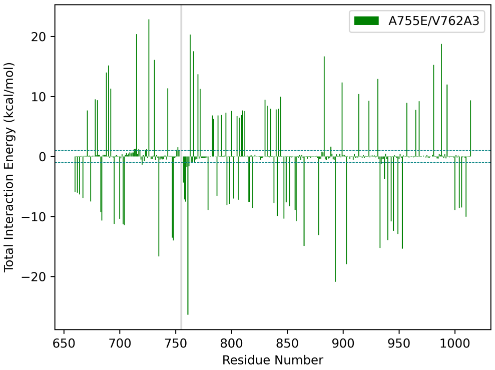


(C) A755E/V762A3

Figure S20: Total interaction energy (Coulomb and van der Waals) with respect to residue 755 (highlighted in gray) in A755E/V762A.

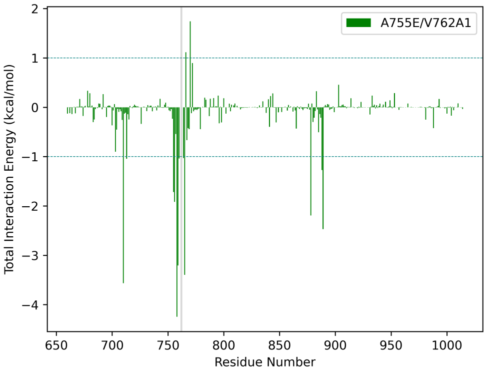


(A) A755E/V762A1

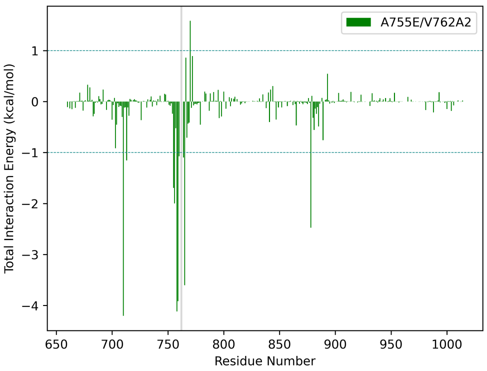


(B) A755E/V762A2

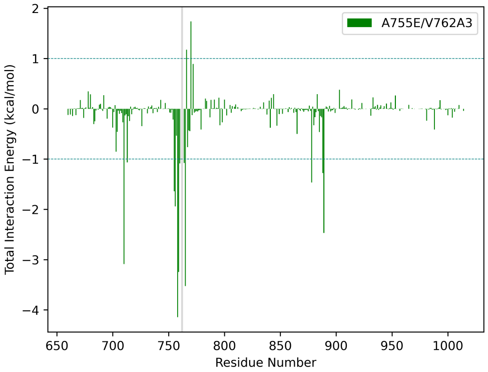


(C) A755E/V762A3

Figure S21: Total interaction energy (Coulomb and van der Waals) with respect to residue 762 (highlighted in gray) in A755E/V762A.

## 4.7. A755L/V762A

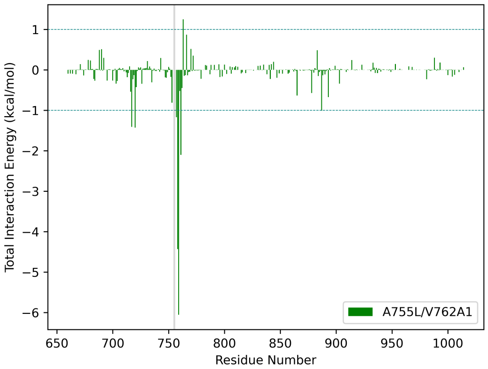


(A) A755L/V762A1

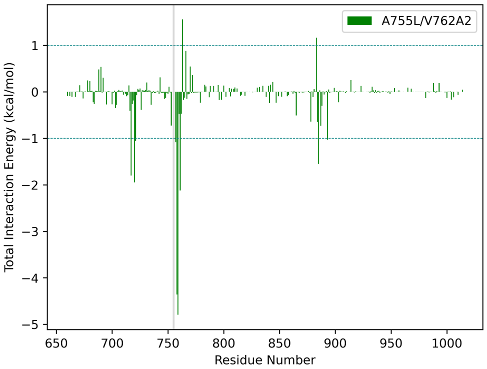


(B) A755L/V762A2

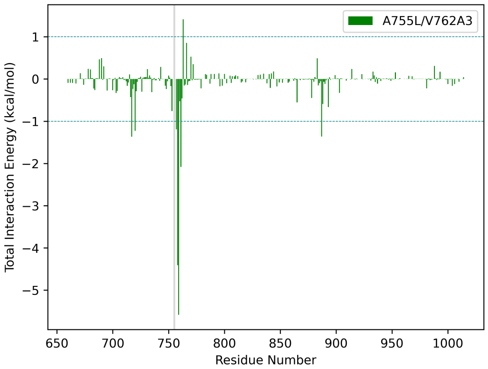


(C) A755L/V762A3

Figure S22: Total interaction energy (Coulomb and van der Waals) with respect to residue 755 (highlighted in gray) in A755L/V762A.

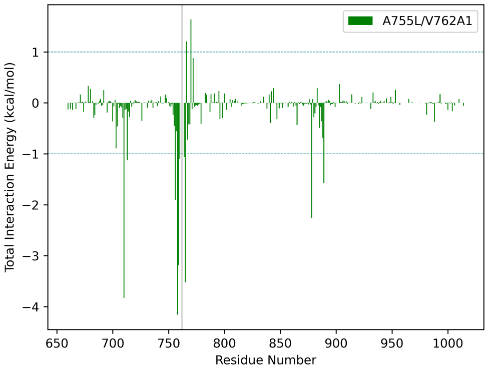


(A) A755L/V762A1

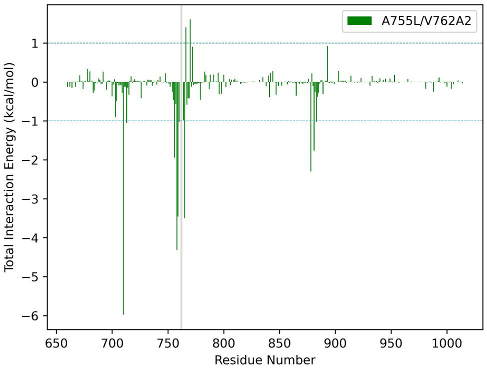


(B) A755L/V762A2

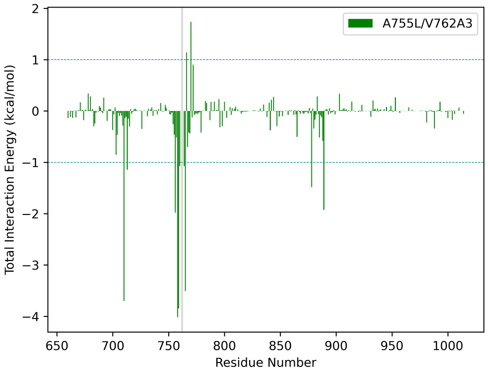


(C) A755L/V762A3

Figure S23: Total interaction energy (Coulomb and van der Waals) with respect to residue 762 (highlighted in gray) in A755L/V762A.

#### 4.8. Difference EDA

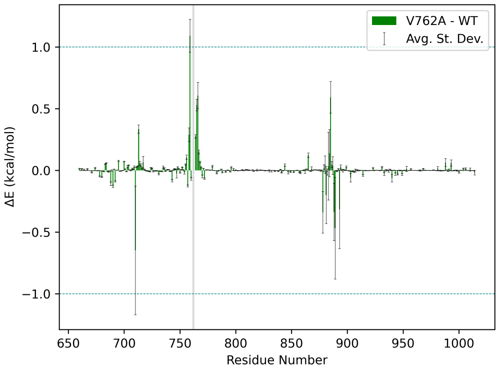


(A) V762A – WT

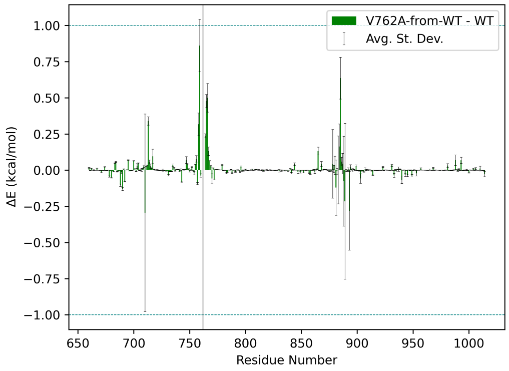


(B) V762A-from-WT – WT

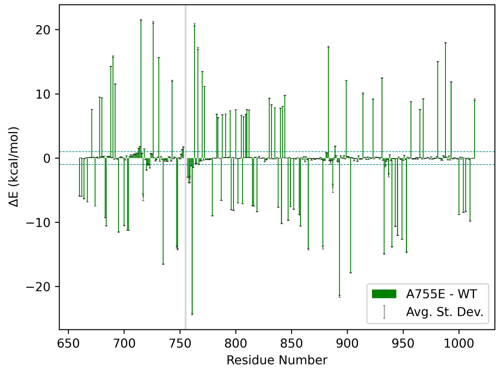


(C) A755E – WT

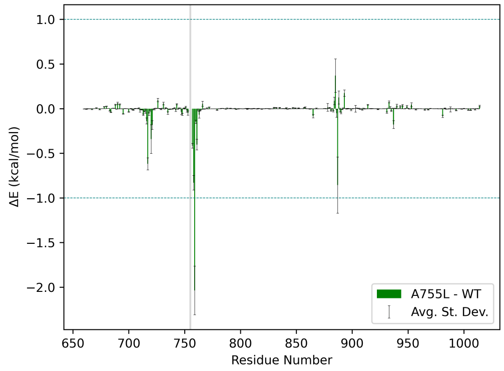


(D) A755L – WT

Figure S24: Difference in average total interaction energy (Coulomb and van der Waals) for the system for the variant – WT. V762A and V762A-from-WT are with respect to the interactions for residue 762, and A755E and A755L are with respect to the interactions for residue 755. The variant position is highlighted gray, and average standard deviation is indicated with black error bars.

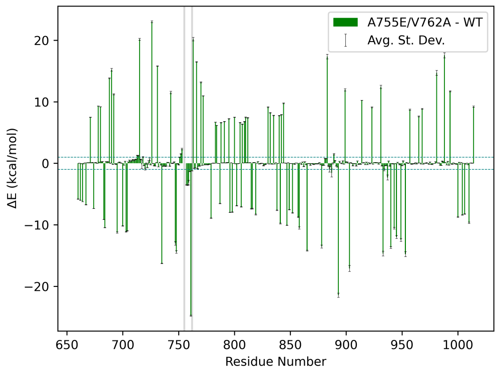


(A) A755E/V762A – WT at residue 755

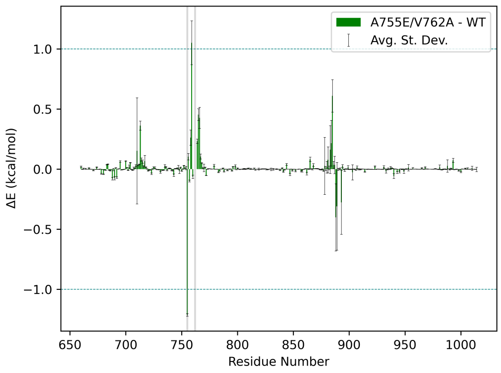


(B) A755E/V762A – WT at residue 762

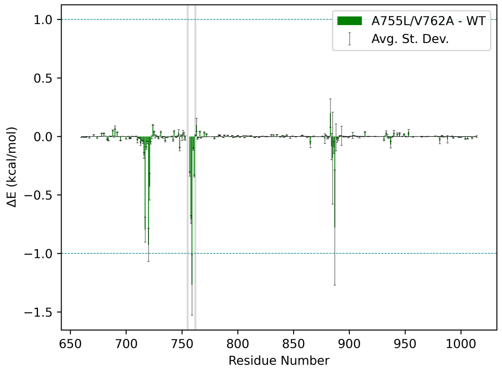


(C) A755L/V762A – WT at residue 755

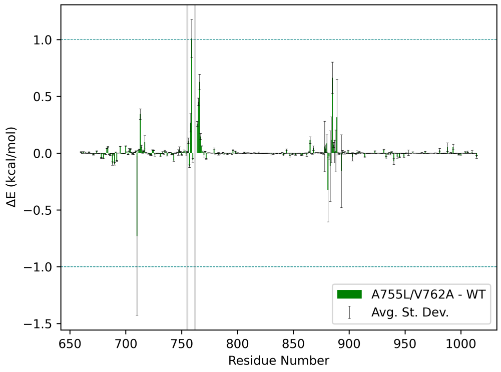


(D) A755L/V762A – WT at residue 762

Figure S25: Difference in average total interaction energy (Coulomb and van der Waals) for the system for the double variant – WT. Each are with respect to the residue specified in the subcaption. The variant positions are in highlighted gray, and average standard deviation is indicated with black error bars.

#### 5. Normal Mode Analysis

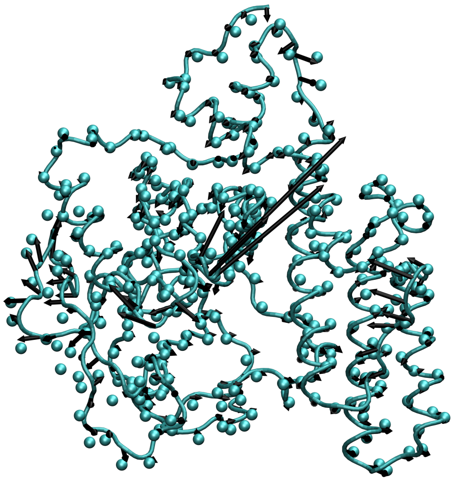

(A) WT

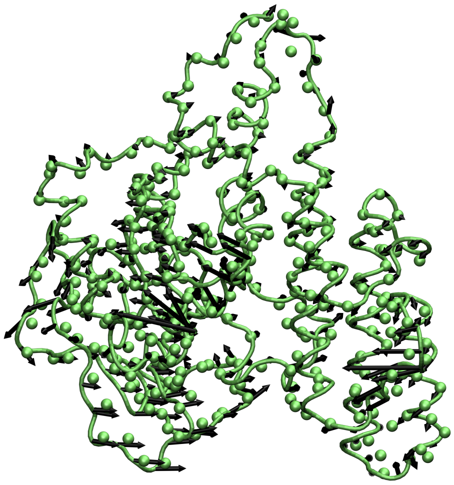

(B) V762A

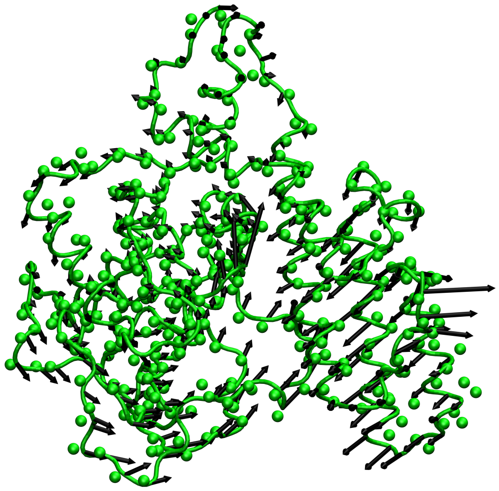

(C) V762A-from-W

Figure S26: First normal mode for a representative simulation of each system.

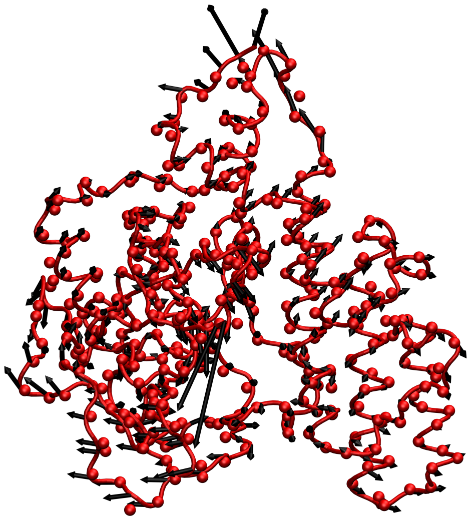

(A) A755E

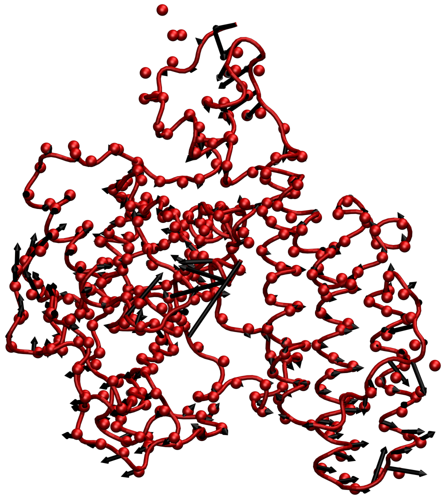

(B) A755E/V762A

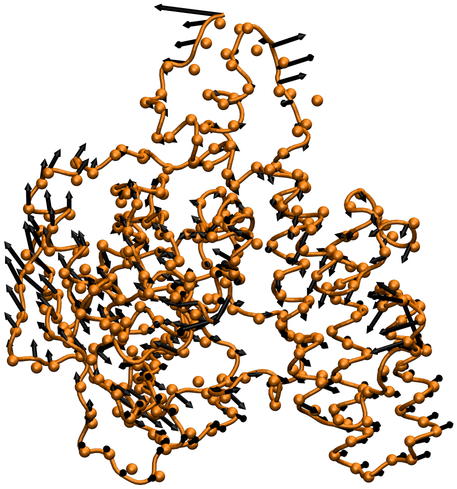

(C) A755L

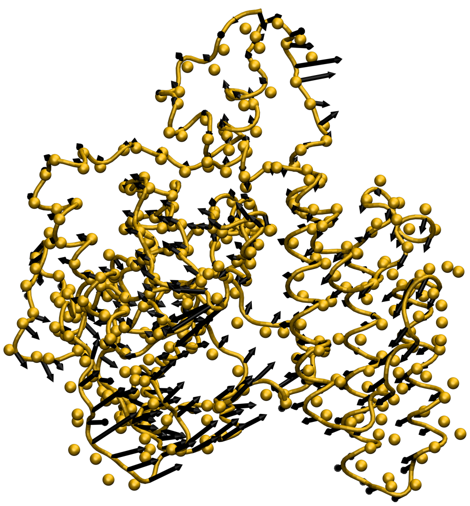

(D) A755L/V762A

Figure S27: First normal mode for a representative simulation of each system.
